# MORGaN: self-supervised multi-relational graph learning for drug target discovery

**DOI:** 10.1101/2025.09.10.675402

**Authors:** Martina Occhetta, Mani Mudaliar, Anniek Myatt, Conrad Bessant

## Abstract

Identifying therapeutically tractable targets remains difficult, partly because disease biology is distributed across multiple molecular layers and relation types, while labeled data are scarce. We present MORGaN, a self-supervised framework for node classification on multi-omic, multi-relation gene networks that learns structure-aware embeddings and outputs calibrated scores to prioritize therapeutic targets. On a pan-cancer graph integrating TCGA multi-omics and diverse biological relation types, MORGaN outperforms state-of-the-art biological node classification models across metrics (AUPR: 0.815 → 0.888; +9%). Ablation studies highlight that both relation diversity and the in-layer fusion architecture are necessary for these gains. Prioritized targets are biologically coherent: high-confidence hits are enriched for pharmaceutically tractable families and ligand–receptor signaling cascades. Post hoc explainability analyses recover compact, pathway-consistent motifs around both known and putative novel targets, and concordance with external resources further supports plausibility. MORGaN thus delivers label-efficient, interpretable node classification for target discovery and can be readily adapted to other diseases, other species, and other node classification tasks. Code and documentation are available at [this link](https://github.com/martina-occhetta/MORGaN).

## 1 Introduction

Drug discovery is complex and time-consuming: bringing a new drug to market can take over a decade and cost upwards of 2.6 billion USD, with failure rates remaining high across all stages of development [1–3]. Drug target identification is a crucial bottleneck: selecting targets whose modulation translates into clinical benefit determines downstream efficacy, attrition, and cost [2–4]. Notably, putative targets with human genetic support are more likely to lead to clinical success, underscoring the value of methods that integrate disease-relevant molecular evidence [5, 6]. In oncology, the challenge is magnified by tumor heterogeneity and context-dependent interactions across genetic, epigenetic, and proteomic layers [7–12].

Conventional methods for drug target discovery often rely on single modalities or curated pathways, risking the loss of cross-layer dependencies and disease-context specificity. Network-based approaches address this in part by modeling protein–protein interactions (PPIs), yet most rely on this single relation type and overlook complementary links such as co-expression, pathway co-occurrence, sequence or domain similarity [9–14]. Capturing systems-level context therefore requires models that integrate diverse molecular evidence and the multiple ways genes relate to one another.

Graph neural networks (GNNs) are well suited to this setting: they combine node features with network topology via message passing to learn context-aware representations [15–17]. Relational Graph Convolutional Networks (RGCNs) extend GNNs to heterogeneous graphs with multiple edge types, aligning with the reality that genes are linked by distinct biological relations [18, 19]. Two practical barriers, however, limit their impact on drug target identification: the scarcity and bias of labeled targets, and the computational cost of naïve multi-relation message passing on large, dense graphs [11, 14].

To address these challenges, we present **MORGaN**, a self-supervised, multi-relational graph learning framework for drug target identification (Fig. 1). MORGaN integrates multi-omic features with six biologically meaningful relation types in a single model. Critically, node features confer disease specificity: cancer-type–resolved multi-omic profiles anchor each gene in its disease context, while the relations provide reusable wiring priors that capture general biological connectivity. During pre-training, a masked autoencoder extends feature-reconstruction objectives [20] to *multi-omic, multi-relation* gene graphs, enabling representation learning from all genes – not only from the small labeled subset – while preserving relation semantics. For scalability, a lightweight relational kernel vertically stacks sparse adjacency matrices and employs basis decomposition to collapse relation-specific message passing into a single sparse–dense operation, substantially reducing per-epoch time (by approximately 80% in our benchmarks) without sacrificing accuracy. Importantly, MORGaN yields mechanistically interpretable outputs: subgraph explanations highlight the minimal gene modules and relation types that support each prediction, situating candidates within pathways and complexes and providing falsifiable hypotheses for experimental follow-up.

**Figure 1:**
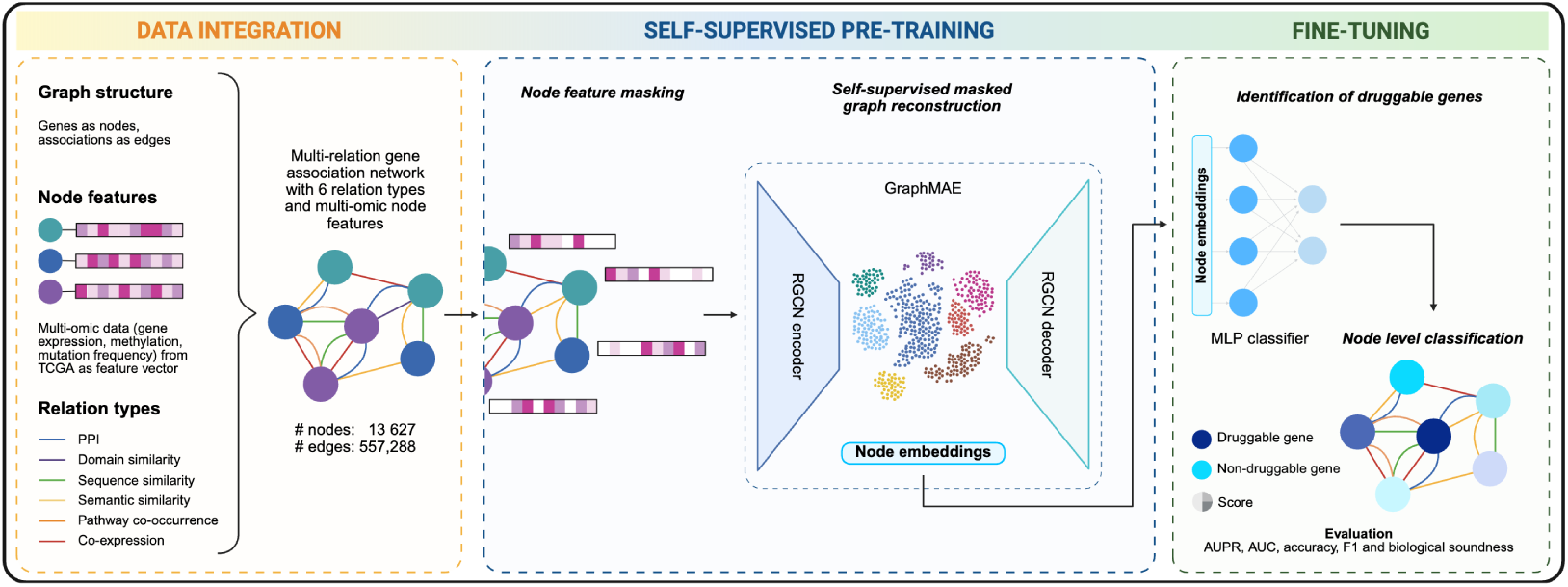
MORGaN overview. *Graph construction and data integration*: build a six-relation, multi-omic gene network. *Self-supervised pre-training*: a RGCN-based masked autoencoder (GraphMAE) reconstructs the missing features and generates node embeddings. *Fine-tuning*: an MLP uses these embeddings to rank drug targets, evaluated with AUPR, AUROC, accuracy and F1.

MORGaN builds on and unifies threads of prior work. SMG applies masked reconstruction to PPIs for cancer and essential-gene prediction under scarce labels [14]; MODIG [21] and MDMNI-DGD [22] extend from single PPIs to five- and six-edge-type multiplex graphs but train relations separately. In contrast, MORGaN incorporates six relation types within an efficient relational encoder and uses self-supervised pre-training to exploit unlabeled biology at scale. Moreover, MORGaN remains disease-agnostic: re-targeting to new contexts requires only substituting features and labels.

By coupling multi-omics integration, relation-aware message passing, and self-supervised learning in a single framework, MORGaN advances graph-based methodology while delivering biologically interpretable results that are directly actionable for systems-level target discovery.

## 2 Results

We evaluate MORGaN on a pan-cancer gene network that couples TCGA multi-omic features (copy-number, expression, mutation, methylation) with six complementary relation types (PPI, co-expression, GO semantic similarity, pathway co-occurrence, sequence, and domain similarity). The model is pre-trained via masked feature reconstruction and then fine-tuned to predict drug targets using stratified train/validation/test splits. We first establish predictive performance and efficiency, and then examine what MORGaN learns biologically – how genes are organized in the latent space, which families and pathways are prioritized, and how predictions sit in the context of the network.

### 2.1 MORGaN consistently outperforms current state-of-the-art

Across six stratified shuffle–split runs, MORGaN delivers the best node classification performance on every metric (Fig. 2; Supplementary Table 3).MORGaN reaches AUPR 0.888 ± 0.004, exceeding the strongest published heterogeneous models (MDMNI-DGD) by +0.073 absolute (+9%). Discrimination likewise improves: AUROC 0.907 ± 0.005 versus 0.877 for MDMNI-DGD (+0.030), and balanced classification follows suit with F1 0.917 ± 0.004 and Accuracy 0.915 ± 0.005. Variability is small across repeats, indicating stable gains rather than favorable splits. All models are evaluated on identical train/validation/test splits with the same node features; heterogeneous models also use the same six-relation graph, isolating the contribution of MORGaN’s architecture and training.

**Figure 2:**
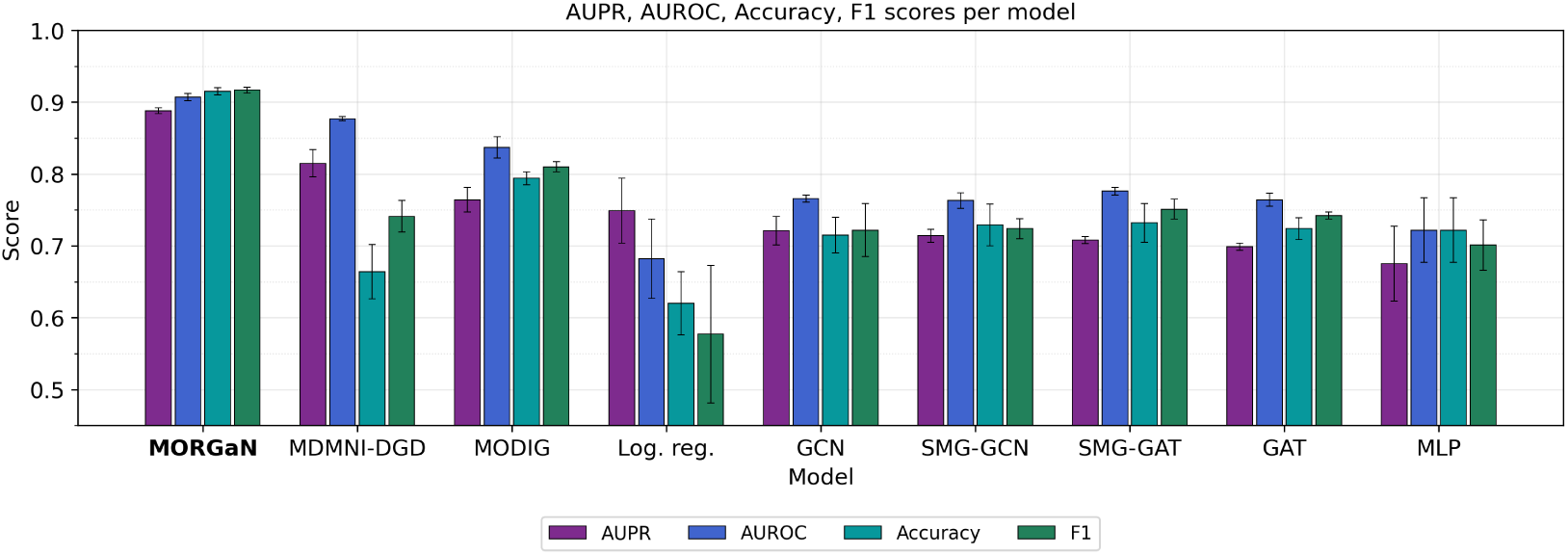
Grouped bars show mean test set performance (AUPR, AUROC, accuracy and F1 score) across the same six stratified shuffle–split runs; error bars denote standard deviation. All models receive the same multi-omic node features; heterogeneous methods (MDMNI-DGD, MODIG) also share the identical six-relation graph. A full description of the models is provided in Appendix C.

A head-to-head comparison within each model family suggests that heterogeneous edges matter, but architecture matters more. Indeed, substituting the single-edge view of GAT with the full six-relation interactome already yields a strong lift (AUPR +0.065 from GAT to MODIG). This gain confirms that signals are not confined to one molecular relationship but are dispersed across many. MORGaN goes further by fusing relations *within* each message-passing layer, allowing information to flow *between* relations as representations are updated. This in-layer cross-talk yields an additional +0.073 AUPR over the strongest heterogeneous baseline (MDMNI-DGD) and leaves even the ablated MORGaN (no pre-training) far ahead. MORGaN seems to capture the complexity of biological systems in a way that traditional models don’t – recovering pathway interplay and complementary cues that architectures processing relations in isolation tend to miss.

Together, these observations show that (i) embracing the full diversity of biological interactions and (ii) employing an architecture specifically designed to fuse those interactions in-layer are both necessary – and mutually reinforcing – for state-of-the-art drug target identification.

We further test robustness under both distribution and task shift: (i) disease shift by re-training MORGaN on a graph with the same relations and Alzheimer’s disease (AD)-specific node features; and (ii) task shift, by applying the framework to essential-gene prediction. We observe qualitatively consistent trends under both types of shift; see Appendix L for full protocols and results.

### 2.2 MORGaN prioritizes a large, biologically coherent set of candidate targets

MORGaN correctly retrieves hallmark cancer drug targets such as *EGFR*, *HER2*, *BRAF*, *ALK*, *MET*, and *RET*. To quantify confidence across all genes, we use a consensus probability score_mean (the mean predicted probability across seeds/splits) and bin genes into tiers: below ≤ 0.60, exploratory (0.60, 0.75], medium (0.75, 0.90], high (0.90, 1.00], plus an additional **top** label for score_mean= 1.0. Using this scheme, MORGaN nominates 1141 *high-confidence* positives and 312 *top* hits, including 163 novel candidates. These predictions are not diffuse; they cluster in pharmacologically tractable families and converge on cancer and immune signaling programs (below). Moreover, an external concordance analysis shows that the majority of high-confidence predictions are independently supported by DGIdb and/or the Finan druggable-genome atlas, with a large three-way intersection (Appendix J, Table 16).

#### 2.2.1 Embeddings organize genes into pathway-coherent clusters

To assess whether MORGaN’s representations organize genes into biologically coherent modules, we projected the learned embeddings with UMAP (cosine; *n*_neighbors_ = 30) and overlaid the predictions. The 2-D map reveals a broad crescent-shaped manifold on which predicted probabilities vary smoothly (Fig. 3a); the majority of high-confidence positives occupy the arc, whereas predicted negatives form a separate island. Crucially, “unlabeled → predicted+” genes (our putative novel targets) co-localize with known positives rather than with negatives (Fig. 3b). A simple *k*-means partition of the UMAP (*k* = 12) further divides the space into contiguous segments (Supplementary Fig. 14); cluster-level enrichment points to distinct biological programs. For example, the bottom left-hand island (cluster6) is enriched for nuclear-receptor and sterol biology together with drug-metabolism modules (Cytochrome P450; Phase I functionalization) and GPCR ligand binding (Supplementary Fig. 15b), consistent with a detoxification/metabolic hub. An adjacent island (cluster3) highlights stress-response and metabolic control (TP53-regulated metabolic genes, glutathione metabolism, mTOR signaling; Supplementary Fig. 15a), while a segment on the outer arc (cluster11) is enriched for receptor-proximal trafficking and signal transduction (retrograde Golgi transport, RHO GTPase cycle, BMP signaling, and PI3K cascade downstream of FGFR2; Supplementary Fig. 15d). Global projections that regress individual drivers onto the centered UMAP coordinates identify modest but consistent axes aligned with the crescent (per-feature *R*^2^ ∽ 0.01 − 0.02), including cancer–type gene-expression contrasts (e.g., BRCA, LUAD, PRAD, LUSC, KIRC/KIRP) and a clustering-based summary (Supplementary Fig. 16). Together, these analyzes indicate that MORGaN arranges genes along interpretable gradients – from metabolic/detoxification and nuclear-receptor programs through stress and growth-factor signaling to GPCR/RHO-cytoskeletal modules – and that newly prioritized, previously unlabeled genes fall into the same neighborhoods as established targets.

**Figure 3:**
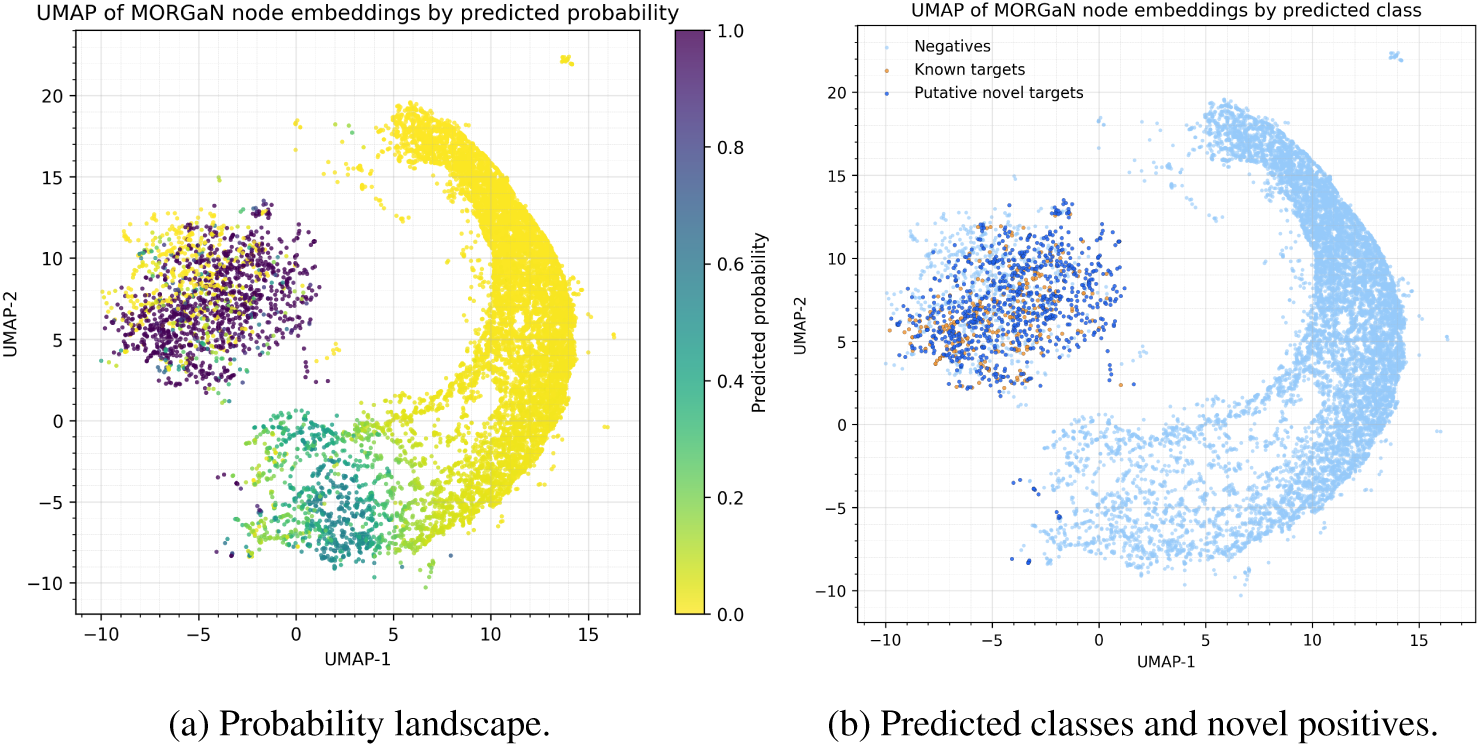
(a) 2-D UMAP of MORGaN embeddings (cosine, *n*_neighbors_ = 30); each point is a gene colored by predicted probability *p*(positive). Probabilities vary smoothly along a crescent-shaped manifold; a separate island concentrates predicted negatives. (b) Same UMAP with layers: background predicted negatives (light), known targets (true positives, small orange points), and novel putative targets (“previously unlabeled → predicted positive” genes, blue). Novel positives co-localize with known positives rather than the negative island, consistent with pathway-coherent prioritization.

#### 2.2.2 Pharmaceutically privileged families are enriched among high-confidence predictions

Across prediction tiers, MORGaN progressively concentrates signal in pharmaceutically tractable classes (Fig.4a). The fraction of GPCRs, ion channels and kinases rises from the “below”/“exploratory” sets to the “high” and “top” tiers, while the residual “other” category shrinks, indicating that higher model confidence coincides with established drug targets [23]. Formal over-representation testing in the top tier (Supplementary Table 13) confirms significant enrichment of GPCRs, ion channels, broad receptors, and cytokine receptors. By contrast, kinases show no enrichment versus the genomic background, consistent with their high baseline prevalence and suggesting that MORGaN shifts attention toward non-kinase, receptor-mediated opportunities. Together, these patterns indicate that MORGaN’s highest-confidence calls naturally align with well-validated target families – particularly GPCR and ion-channel biology – while still leaving room for diverse mechanisms in the remaining fraction, which we examine in downstream pathway and network analyses.

Immune-relevant families follow the same pattern: among high and top calls we observe substantial representation of cytokine signaling (*n* = 42 and 17), chemokine axes (*n* = 16 and 9), checkpoint/co-stimulatory molecules (*n* = 14 and 8), and antigen-presentation components (*n* = 6 and 3), aligning with the cytokine-receptor enrichment and pointing to both tumour-intrinsic and microenvironmental immune levers.

#### 2.2.3 Pathway-enriched programs concentrate in GPCR and RTK–PI3K/ERK signaling

We assessed pathway over-representation on the High tier using a Fisher exact test against the genome-wide background with Benjamini–Hochberg correction. High-confidence predictions were strongly enriched for ligand–receptor signaling and growth-factor cascades (Fig. 4b). The most prominent signal was Neuroactive ligand–receptor interaction (odds ratio, OR ≈ 18 − 20; *q <* 10^−50^), accompanied by Reactome GPCR modules – Signaling by GPCR, GPCR ligand binding, and Class A/Rhodopsin-like receptors (OR ≈ 5 − 8; *q <* 10^−30^). In parallel, we observed pronounced enrichment for growth-factor receptor tyrosine kinase pathways including RTK → RAS → ERK and RTK → PI3K branches (OR ≈ 20 − 26; *q <* 10^−20^), together with PI3K–AKT signaling. A generic Cancer pathways term also appears, but the signal is mechanistically concentrated in GPCR/RTK axes. These results align with the family-level tractability analysis – where GPCRs, ion channels, and receptors are over-represented – and indicate that MORGaN preferentially elevates targets embedded in well-studied signaling circuits with clear translational leverage.

**Figure 4:**
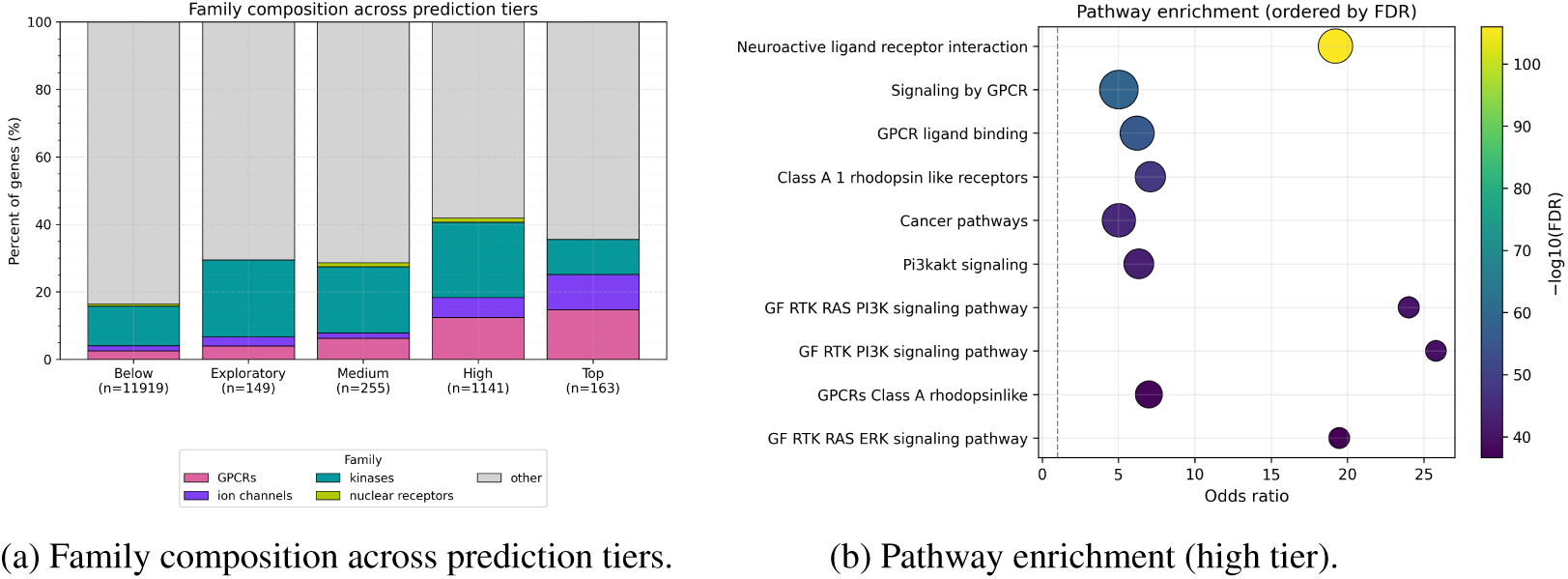
(a) Stacked bars show the fraction of genes in pharmaceutically privileged families – GPCRs, ion channels, kinases, nuclear receptors – versus “other” for each prediction tier. The proportion of tractable classes increases from Below/Exploratory to High/Top, while the “other” fraction shrinks, indicating that higher model confidence concentrates in druggable modalities. (b) Bubble plot of over-representation analysis for the High tier (Fisher’s exact test versus genome-wide background; Benjamini–Hochberg FDR). Bubble size encodes the number of overlapping genes; color encodes − log_10_(FDR); dashed line marks odds ratio = 1.

#### 2.2.4 Network context: MORGaN is not just a “hub detector”

To understand how predicted targets are positioned in the interactome, we examined degree, betweenness, and clustering. Top hits occupy more central and cluster-forming positions than background genes (all *p <* 10^2^), but correlation analyses (Supplementary Fig. 11) show that MORGaN is not simply a hub detector. Across all genes, the correlation between score and network-derived features is weak: degree *ρ* = 0.25 (partial 0.04), betweenness *ρ* = 0.30 (partial 0.17), and clustering *ρ* = 0.11 (partial 0.07); running-median trends indicate high scorers span a wide degree range but are modestly enriched along information-flow routes and within local modules. Biologically, this pattern is desirable: pure hubs can be toxic or essential, whereas genes that broker flow between modules or sit in compact neighborhoods often mediate tractable, disease-relevant processes. These analyses therefore support a systems-level notion of targetability aligned with pathway wiring rather than raw connectedness.

### 2.3 Local explanations recover compact, pathway-coherent evidence per gene

To inspect *why* MORGaN assigns high confidence to specific genes, we applied GNNExplainer to the trained model (methods in Appendix I). The explainer learns soft masks over edges and feature dimensions, yielding (i) a small *explanation subgraph* per gene and (ii) a cancer-type × omic feature–importance profile. Across examples, the selected subgraphs are compact (visualized as the top-20 edges) and map onto recognizable signaling motifs, while the feature masks highlight disease contexts and modalities that plausibly support the call (Fig. 10, 9).

Snapshots illustrate three recurring patterns. (i) Receptor–checkpoint crosstalk around known targets. For *EGFR*, salient neighbors include *TP53*, *CDK2*, and *CTNNB1*, tying RTK signaling to cell-cycle and Wnt control; features concentrate in lung cancer CNAs and expression, matching clinical use. (ii) Multi-hop pathway context. For *NOTCH1*, the explainer emphasizes an ERBB4→MAPK9 route and the *RBPJ* transcriptional module, indicating that MORGaN leverages how Notch feeds into MAPK and downstream transcription rather than counting direct interactors. (iii) Mechanistic neighborhoods for novel candidates. For *LAMA3*, the subgraph links integrins and SMADs (ECM–integrin–TGF*β* crosstalk); for *IL4R*, edges to *AKT2*, *RAC1*, and *TP53BP1* capture immune-to-survival and cytoskeletal routes. In both cases, feature heatmaps point to tissue contexts (e.g., bladder/thyroid for *LAMA3*, colorectal/lung for *IL4R*) and modalities (expression, CNA) that align with the network evidence.

Together, these local explanations indicate that high MORGaN scores are supported by structured network motifs and relevant omic signals, not by degree alone – complementing our global centrality analyses – and they translate directly into testable, pathway-level hypotheses for experimental follow-up (Appendix I).

#### 2.3.1 Genetic dependency signals corroborate predictions and reveal lineage-specific vulnerabilities

Cross-referencing DepMap reveals that all novel putative targets have at least one supporting dependency signal, and we identify candidates with lineage-preferential dependencies – precisely the kind of context that enables rational stratification. Examples include SMG1 (biliary tract), KRAS (pancreas), and cohesin/replication-associated factors (FANCM, DSCC1, CHTF18) in fibroblast-like contexts, as well as CHP1, CDK6, and ERBB4 in adrenal lineage. Coupled with family and pathway annotations, these data guide concrete experimental programs: for instance, focusing GPCR modulators on specific lineages where genetic dependency is strongest, or combining RTK inhibitors with downstream PI3K/mTOR agents in contexts supported by both network placement and dependency evidence.

#### 2.3.2 Indication-focused case studies of novel MORGaN candidates

To translate aggregate performance into disease-facing hypotheses, we present indication-focused case studies drawn from the top novel putative target set, illustrating how MORGaN’s rankings map to tractable biology and concrete experimental avenues.

#### Non-small-cell lung cancer (NSCLC)

Among MORGaN’s putative novel targets, we highlight EREG, that maps to tractable axes in NSCLC biology. EREG (epiregulin) is an EGFR/ERBB ligand that can establish autocrine signaling and is associated with aggressive behavior in NSCLC: functional work shows EREG drives proliferation and invasion via EGFR/MEK/ERK activation, and high EREG expression portends poorer outcomes in patient cohorts. These data suggest an actionable dependency in EREG-high tumors and motivate testing ligand-axis blockade (e.g., pan-EGFR/ERBB inhibition) in the MORGaN-prioritized subset [24–26].

#### Acute myeloid leukemia (AML)

MORGaN surfaces immune-checkpoint and cytokine-axis genes that are rarely annotated as drug targets in AML labels but have growing translational support. CD274 (PD-L1) is up-regulated on AML blasts and in the leukemic microenvironment, dampening anti-leukemia T-cell activity; multiple studies and reviews now document PD-L1/PD-1 pathway engagement in AML and provide a mechanistic rationale for biomarker-guided checkpoint blockade and combination strategies. MORGaN’s consistent prioritization of CD274 strengthens the case for systematic re-evaluation of PD-(L)1 targeting in molecularly defined AML subsets [27–29]. As a complementary axis, IL2RA (CD25) – predicted as a high – confidence positive—has been implicated in leukemic stem cell programs and adverse biology in myeloid disease, with reports linking IL2RA expression to stemness and immune evasion. These observations, together with existing CD25-directed agents, nominate CD25-high MORGaN candidates for prospective validation and potential therapeutic exploration.

#### Ovarian cancer

MORGaN repeatedly flags GPCRs in the lysophosphatidic-acid (LPA) pathway, including LPAR2, as consistently positive. The LPA–LPAR signaling axis is a well-described driver of ovarian cancer proliferation, migration, and peritoneal dissemination, with LPA present at high levels in malignant ascites and multiple receptor subtypes (including LPAR2) contributing to pro-metastatic phenotypes [30–34]. The convergence of MORGaN’s predictions with this pathway supports testing LPA-receptor antagonism (or pathway-directed combinations) in the MORGaN-enriched subset of ovarian tumors.

### 2.4 MORGaN delivers accurate predictions at a fraction of the computational cost

On identical hardware and training budgets, MORGaN attains state-of-the-art accuracy with markedly lower wall-clock time. A complete pre-train → fine-tune cycle takes 24.3±2.9 s – about 65 and 23 faster than MODIG and MDMNI-DGD, respectively (Supplementary Table 2). The speedup arises from two design choices in the encoder: vertically stacked *sparse* message passing that fuses all relations within each layer, and basis-matrix weight decomposition that reduces per-relation parameters without sacrificing expressivity (see §4.2). Full hardware specification, timing protocol, and per-stage breakdown are provided in Appendix D.

### 2.5 Ablations confirm MORGaN’s dependence on real biology

To dissect which inputs and design choices drive performance, we ran three ablations on the cancer graph, each repeated over the same six stratified shuffle–split runs (reporting mean ± s.d.). We quantify changes as ΔAUPR relative to the full six-relation, four-omics model (details in Appendix F).

#### Drop-one edge type

Starting from the full six-relation graph (PPI, pathway co-occurrence, GO semantic similarity, co-expression, sequence similarity, domain similarity), we removed one relation at a time and re-trained the pipeline. Fig. 5 (blue) shows the resulting AUPR distributions; the line at *n* = 6 is the unablated reference. The largest loss arises when removing *GO semantic similarity* (ΔAUPR 0.010; 0.888 0.878), with PPI and pathway co-occurrence nearly as impactful (each 0.009). Biologically, this pattern is intuitive: GO similarity encodes *functional proximity* across processes and cellular components, capturing long-range dependencies (e.g., genes that never touch but act in the same complex program); PPI edges represent physical wiring within complexes; and pathway co-occurrence reflects coordinated participation in signaling cascades. Together, these relations describe complementary aspects of the same system. By contrast, dropping sequence similarity or co-expression slightly improves AUPR (to 0.900 and 0.899), suggesting redundancy/noise in those layers. Importantly, every five-relation variant still outperforms the best single-relation baseline (GO-only, 0.883), indicating that the signal is distributed across heterogeneous biological relationships rather than concentrated in any one layer. Consistent with the edge ablations, averaging over all *n*-relation subsets shows monotonic gains that saturate by *n* = 5–6 (Supplementary Fig. 13), reinforcing the biological premise that drug-relevant mechanisms are multiplex and that MORGaN’s in-layer fusion captures that reality.

**Figure 5:**
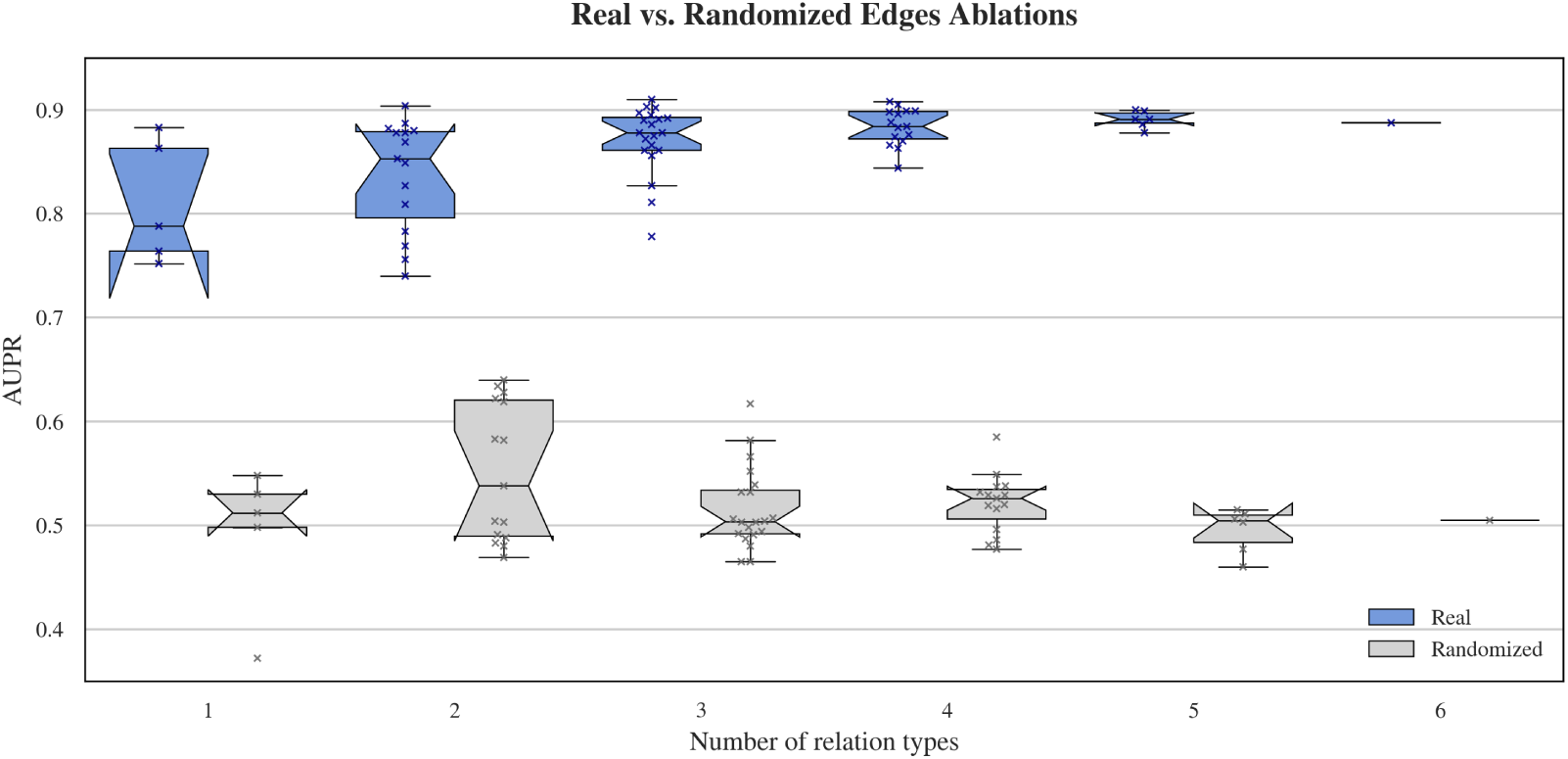
For every subset size *n* we plot AUPR over all combinations of real edges (blue) and their randomized counterparts (grey). The “box” at *n* = 6 reduces to a thin line because only one configuration – the full graph – exists.

#### Degree-preserving randomized controls

To test whether gains reflect genuine biology rather than additional parameters or edge density, we degree-preserving–shuffled each relation and repeated the analysis (grey boxes in Fig. 5). AUPR collapsed toward chance (~0.5) across all subset sizes, demonstrating that MORGaN’s accuracy depends on real network topology and cross-relation organization, not mere edge density.

#### Leave-one-omics-out

A similar analysis on node features shows that *copy-number alterations (CNA)* carry the strongest standalone signal (highest single-modality AUPR), whereas bulk *gene expression* is the noisiest—its removal increases AUPR by +0.009, aligning with prior evidence that high-dimensional expression with limited samples can dilute signal. Nonetheless, the *full four-omics* model yields the best overall trade-off, achieving the top AUROC (0.907) and near-top AUPR by recovering false negatives missed by CNA alone (Appendix F.2).

Together, these ablations show that (i) long-range functional context captured by GO terms, alongside PPI and pathway co-occurrence, is essential; (ii) the multi-relation structure is informative – scrambling biology erases the gains; and (iii) complementary omics improve discrimination despite noisy modalities. Robustness to alternative PPI sources holds (STRING performs best, but MORGaN remains superior across interactomes; Appendix Table 4).

### 2.6 MORGaN is robust to class imbalance

We next quantified the robustness to the class balance of the training set. We swept the negative:positive ratio used (0.25, 0.5, 1.0) while keeping the training settings fixed. Across six repeated splits, MORGaN sustained uniformly high precision–recall, with only a modest decline as negatives increased (AUPR ≈ 0.98 → 0.91; Supplementary Fig.12, left). AUROC remained stable and near-maximal (peaking around the 1:2 balance and decreasing slightly at 1:1; Supplementary Fig.12, middle), while Accuracy varied little (Supplementary Fig.12, right), consistent with its known sensitivity to class prevalence. Early-stopping checkpoints were typically as good as, or slightly better than, end-of-training models at the more imbalanced settings, suggesting a small regularization benefit without altering conclusions. Overall, these trends indicate that MORGaN’s ranking of putative targets is robust to plausible shifts in label prevalence, which is critical when deploying models across disease areas.

## 3 Discussion

MORGaN is built on the idea that integrating weak, heterogeneous signals improves target discovery. By coupling multi-omic features with a multi-relation interactome and fusing relations within each layer, it consistently outperforms strong baselines while remaining fast enough for iterative exploration. Sensitivity analyses show these gains persist under plausible shifts in class balance, and ablations indicate they depend on genuine network structure rather than edge density or parameter count. Overall, the results support that target signal is distributed across complementary biological relationships and can be captured by relation-aware message passing grounded in multi-omic context.

### Biological coherence

High-confidence predictions are enriched in pharmaceutically tractable families (GPCRs, ion channels, receptor classes) and concentrate in pathways with clear translational leverage (GPCR signaling and RTK→PI3K/ERK cascades). In the learned representation, genes arrange along interpretable gradients; previously unlabeled positives co-localize with known targets and fall into pathway-enriched clusters. Network analyses further show that MORGaN is not merely recovering hubs: scores correlate only weakly with degree, betweenness, and local clustering, highlighting candidates on information-flow routes and within coherent modules. Brief indication-focused case studies illustrate how these signals translate into testable hypotheses.

### Why does MORGaN work?

Two features appear crucial to MORGaN’s performance. First, relations are fused *within* each message-passing layer, allowing information from different relations to mix during propagation rather than being processed in isolation. Second, masked multi-omic pre-training provides a strong prior when labeled positives are scarce, improving data efficiency at fine-tuning. Edge-type ablations (GO/PPI/pathway co-occurrence) and leave-one-omics-out results (CNA strongest alone; other modalities recover missed cases) align with this view and suggest design principles for future graphs and models.

### Limitations

However, some limitations remain. Labels for drug targets are incomplete and potentially biased toward well-studied families, and our bulk multi-omic features may dilute context-specific signals present at single-cell resolution. The interactome is also incomplete and uneven across sources; although results are robust across alternative PPI layers, any fixed graph can miss disease-specific wiring. Finally, we do not address downstream concerns such as modality safety or on-target toxicity; prioritization as a therapeutic target based on its mechanistic role in pathways is necessary but not sufficient for “clinically actionable”.

### Future work

We see three immediate avenues to increase translational value. (i) Context specificity: MORGaN already surfaces lineage-preferential dependencies; focusing training and evaluation within a cancer type (or molecular subtype) should sharpen this signal and enable biomarker-guided stratification. Incorporating cell-type–resolved and perturbational readouts (e.g., CRISPR/perturb-seq) could further refine context. (ii) Safety and tractability priors: penalizing network centrality or integrating orthogonal toxicity proxies (e.g., intolerance metrics, essentiality screens, or tissue expression breadth) would convert MORGaN’s single-objective ranker into a multi-objective optimizer over efficacy and risk. (iii) Chemistry and structure: adding ligandability features (pocket descriptors, AlphaFold-derived site annotations, docking/electrostatics summaries) would connect biological coherence to physical plausibility.

Our framework delivers accurate, fast, and interpretable rankings with compact, pathway-consistent local rationales suitable for experimental follow-up. Disease specificity is encoded in node features, biological wiring is captured by the relation layers, and task definition is set by the labels, allowing the same relation-aware encoder to be redirected to indication-specific graphs, cross-species networks, or other node-level problems without re-engineering. In practice, MORGaN provides a direct route from heterogeneous molecular data to mechanistically grounded target hypotheses, and a reusable scaffold the research community can adapt across contexts.

## 4 Methods

### 4.1 Graph construction and data integration

We represent the gene interaction landscape as a heterogeneous, multi-relational graph *G* = (V, E, R), where each node *v_i_* ∈ V represents a gene. For each biological relationship type *r* ∈ R, we define a relation-specific edge set E*_r_* ⊆ V × V and we define the full graph as the union over all relations *ε* = U_*r∈R*_ *ε_r_*. The pan-cancer graph contains 13 627 genes and 557 288 edges across six relation types. For further details, see Appendix A.

#### Relations

Following MDMNI-DGD [22], we incorporate six biologically grounded relation types, based on protein-protein interaction networks (PPI), gene co-expression, pathway co-occurrence, gene ontology semantic similarity, and sequence similarity. Self-loops are added to preserve each gene’s own features during message passing. For further details, see Appendix A.2.1.

#### Node features

Each gene node *v_i_* is associated with a multi-omic feature vector *x_i_*, obtained by concatenating log10-transformed somatic mutation frequencies, copy number alteration (CNA) z-scores, DNA methylation *β*-values, and log-normalized gene expression values. All features are extracted from The Cancer Genome Atlas (TCGA) pan-cancer dataset [14, 35], spanning 29,446 tumor samples across 16 cancer types, as in SMG [14]. For further details, see Appendix A.2.2.

#### Labels

Positive labels correspond to Tier 1 targets defined by Finan et al. [4], i.e. proteins with approved drugs or clinical candidates; an equal number of negatives is randomly sampled from the remaining non-target genes to balance class distributions. See Appendix A.2.3.

### 4.2 Model architecture

We adopt the relational graph convolutional network (RGCN) of Schlichtkrull et al. [18] and convert it into a *graph masked autoencoder* (GraphMAE) [20].

### Message passing formulation

For layer *l* the hidden state of node *v_i_*is updated via:

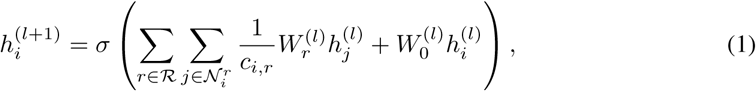

where *h*^(*l*)^_*i*_ represents the hidden state of node *v_i_* at layer *l*, N*^r^*_*i*_ is the set of *i*’s neighbors under relation *r*, *W*_*r*_^(*l*)^ and *W*_0_^(*l*)^ are trainable relation-specific and self-loop weight matrices, respectively, and *c_i,r_* is a normalization constant to ensure numerical stability.

### Vertical stacking for sparse message passing

To exploit the fast sparse–dense multiplication (spmm) available in PyTorch while still updating all relation types at once, we concatenate the *R* relation-specific adjacency matrices {*A_r_*}_*r*=1_^*R*^ *vertically* into a single sparse block matrix *A_v_* ∈ R^(*RN*)×^*^N^*, as introduced by Thanapalasingam et al. [19]. During each RGCN layer, we first mix topological and feature information with one call to spmm(*A_v_, X*), producing a relation-expanded feature matrix of shape (*RN*) × *d_in_*. This matrix is then reshaped back to *N* × (*R d_in_*) and multiplied by a stacked weight matrix to yield the next-layer embeddings. Because the projection to higher dimensions happens *after* the sparse multiplication, vertical stacking keeps memory usage low and scales well to large graphs with modest input dimensionality.

### Weight decomposition

To manage parameter complexity with multiple relation types, we implement basis decomposition [18]. Each relation-specific matrix is expressed as a linear combination of a shared set of *B* basis matrices {*V_b_*}*^B^*_*b*=1_:

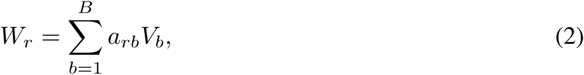

where *V_b_* ∈ R*^d^*^in×^*^d^*^out^ are global basis matrices shared across all relations, and *a_rb_* ∈ R are relation-specific learnable coefficients. This formulation significantly reduces parameter count compared to using unique weights per relation, while preserving expressiveness through learned compositions. In our implementation, we set *B* = 2 to strike a balance between model flexibility and generalization capacity.

### Normalization and dropout

Each layer applies layer normalization to the concatenated relation outputs, adds a residual connection, and then dropout (*p* = 0.2).

### Implementation details

All models are implemented in PyTorch 2.6.0 [36] and PyTorch-Geometric 2.6.1 [37]. Relation weights *W_r_*, bases *V_b_*, and coefficients *a_r,b_*use Xavier uniform (gain = 2 for PReLU) initialization; the self-loop matrix *W*_0_ uses Kaiming initialization. We fix random seeds (Python, NumPy, PyTorch, PyG) to 0 and 1 and report mean ± std over 3 runs per seed.

### 4.3 Training

We adopt a two-phase training strategy with Adam [38].

### Self-supervised pre-training

Following GraphMAE [20], we randomly mask 50% of node features and reconstruct them using a scaled cosine loss:

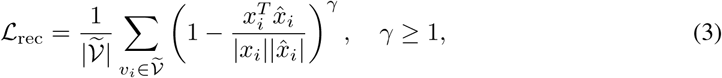

where *x_i_* and *x̂_i_* denote the original and reconstructed feature vectors, respectively, and *γ* controls the loss sharpness. Pre-training runs for 100 epochs with an initial learning rate 10^−2^, weight decay 10^−3^, cosine decay 10^−6^, *γ* = 3, and early stopping (patience 10). Hyper-parameters were selected via a grid sweep (see Appendix B); the best configuration is used throughout the paper. Through this pre-training stage, the model learns compressed embeddings that encode both multi-omic profiles and relational context, serving as a robust foundation for downstream classification.

### Fine-tuning (supervised)

The frozen embeddings feed an MLP classifier optimized with weighted binary cross-entropy:

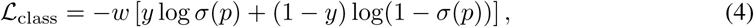

with label-dependent weights *w* to handle class imbalance. We train for up to 200 epochs (learning rate 5 × 10^−3^, weight decay 10^−4^, gradient-clip 1.0) with early stopping (patience 20) on validation AUPR. Hyper-parameters were selected in the same sweep used for pre-training (see Appendix B).

### 4.4 Experimental setup and evaluation

#### Repeated shuffle-split validation

We generate two independent, stratified train/validation/test splits (80% / 10% / 10% of nodes) using different random seeds. Each split is trained three times with different weight initializations, giving six runs in total. We report mean ± s.d. of AUPR, AUROC, Accuracy, and F1 across these runs.

#### Baseline models

We benchmark MORGaN against eight alternatives that span feature-only, homogeneous-graph and heterogeneous-graph approaches:

1. *Logistic Regression* – feature-only
2. *Multilayer Perceptron* (MLP) – feature-only)
3. *GCN* – vanilla graph convolution on a 1-dimensional PPI graph
4. *GAT* – graph attention network on the PPI graph
5. *SMG-GCN* [14] – GCN with self-supervised pre-training on the PPI graph
6. *SMG-GAT* [14] – GAT with self-supervised pre-training on the PPI graph
7. *MODIG* [21] – heterogeneous graph model without pre-training
8. *MDMNI-DGD* [22] – heterogeneous graph model without pre-training

The two *feature-only* models use the concatenated multi-omic vectors. The four *homogeneous* baselines (3-6) operate on a *single-relation* PPI graph and therefore lack the multi-relational context exploited by MORGaN. The two *heterogeneous* baselines (7-8) share the full multi-relational topology with MORGaN but do not include its self-supervised pre-training stage. All models receive identical node features and use the same train/validation/test splits; hyper-parameters are selected by grid search on the validation fold. See Appendix C for further details.

This design cleanly isolates MORGaN’s architectural and training contributions while ensuring a fair, rigorously repeated comparison to both feature-based and graph-based alternatives.

#### Interpretability

For high-confidence predictions (*p >* 0.9), we use GNNExplainer [39] to highlight the network edges and gene features that most drive each call. We then use Enrichr to run *pathway enrichment analysis* – a simple check of whether those highlighted genes occur together in well-known biological pathways more often than expected by chance – reporting pathways that pass a false discovery rate (FDR) threshold of *<* 0.05.

## Data and code availability

The code for MORGaN, including data preprocessing, model training, and evaluation scripts, is available on GitHub. The processed multi-omic feature matrices and relational network adjacency matrices are obtained from publicly accessible resources, including TCGA [35], STRING [40], CPDB [41], IRefIndex [42], MultiNet [43], and PCNet [44]. Instructions for data reconstruction and full reproducibility are available in the repository. The full predictions and the list of novel putative targets is available as supplementary material.

## Author contributions statement

All authors contributed to study design. M.O. implemented the model and performed all analyses.

M.M. and A.M. provided feedback throughout. M.O. and C.B. wrote and revised the manuscript. All authors approved the final version.

## Acknowledgments

This work is supported in part by funds from the UKRI Biotechnology and Biological Sciences Research Council (BB/X51181X/1) and Recursion, as part of the UKRI-BBSRC AI for Drug Discovery Collaborative Training Partnership.

## Appendix

## A Graph construction and data integration

### A.1 Graph summary and descriptive statistics

Table 1 lists the edge count and filtering threshold used for each of the six relation types that form the heterogeneous gene network. The graph is moderately sparse (overall density *<* 0.004), with a heavy-tailed degree distribution typical of biological interaction maps (details in the supplied Jupyter notebook). All subsequent experiments use this exact graph unless stated otherwise.

**Table 1:**
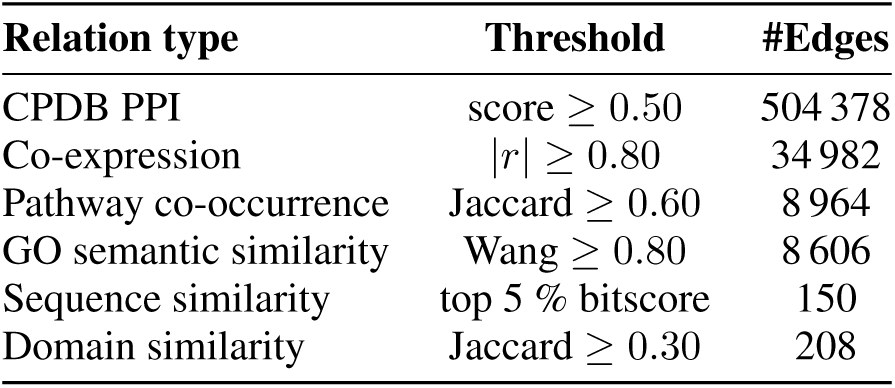
Edge statistics for the heterogeneous gene graph.

### A.2 Components

#### A.2.1 Relations

The heterogeneous MORGaN graph contains six *complementary* edge types. Each captures a different notion of functional similarity; combining them lets the model reconcile noisy, partially overlapping evidence rather than over-focusing on any single assay. We consider the following relation types:

- **Protein–protein interaction (PPI).** Proteins are large biomolecules composed of amino-acid chains encoded by genes. A PPI edge is added when two proteins form a physical complex – e.g. an enzyme binds its substrate or two receptors dimerize – detected by assays such as yeast-two-hybrid or affinity purification. We connect the *genes* that encode the interacting proteins with an undirected edge. Because small molecule drugs also act at this physical level, PPI edges supply high-resolution mechanistic context. High-confidence protein-protein interactions are obtained from one of STRING-db [40] (score ≥ 0.8), CPDB
- [41] (score ≥ 0.5), IRefIndex v.1 and v.4 [42] (score ≥ 0.8), and PCNet [44] (default threshold). CPDB is used as a default.
- **Co-expression.** RNA-seq quantifies how often each gene is transcribed across thousands of samples; higher counts mean the gene is more active. If two genes’ expression profiles are consistently correlated, we add an edge, reflecting shared regulation by common transcription factors or signaling programs – even when their proteins never touch. Co-expression therefore contributes *regulatory* information that PPI alone cannot provide. An edge is added between genes with an absolute Pearson correlation ≥ 0.7 across 79 healthy human tissues, based on GSE1133 [45].
- **Pathway co-occurrence.** KEGG [46, 47] curate step-by-step biochemical pathways (e.g. “MAPK signaling”). Genes that appear in the *same* pathways are linked because they participate in a shared biological process. This injects human knowledge and adds a loose sense of up-/down-stream directionality without exploding the number of edge types. We compute the Jaccard similarity of KEGG [46, 47] pathway memberships:

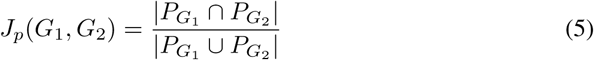

and include an edge where similarity ≥ 0.60.

- **GO semantic similarity.** The Gene Ontology (GO) is a controlled vocabulary with three name-spaces: *Biological Process* (what the gene does), *Molecular Function* (how), and *Cellular Component* (where) [48, 49]. Terms are assigned by curators and automated pipelines.

GO edges generalize “same pathway” and cover genes that lack rich KEGG annotations. We compute the geometric mean of best-match-average (BMA) Wang scores [50] across the GO Biological Process (BP), Molecular Function (MF), and Cellular Component (CC) ontologies [48, 49]:

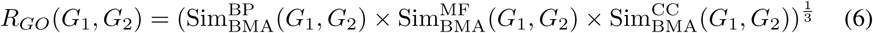

and add an edge where *R_GO_*(*G*_1_*, G*_2_) *>* 0.80.

- **Sequence similarity.**A sequence similarity edge joins proteins whose sequences align with high statistical confidence. Such homology implies a common ancestor and often a shared 3-D fold or catalytic pocket, allowing MORGaN to transfer knowledge from well-studied family members to poorly characterized relatives. We add an edge to the top 5% BLAST bit-scores (normalized for sequence length) between non-identical gene pairs.
- **Domain similarity.** Pfam domains are recurrent, modular sequence blocks that fold into functional units (e.g. SH2, zinc-finger). We connect two proteins if the Jaccard similarity between their Pfam domain sets exceeds 0.30 [51]. Whereas full-length sequence similarity is global, domain similarity edges focus on the local pockets – pinpointing druggable pockets that recur across otherwise dissimilar proteins, which has proved useful for scaffold hopping in medicinal chemistry.

**Why multiple relations?** Biology is inherently multi-scale: genes can be co-expressed yet never touch, or interact directly yet be regulated in opposite ways. Integrating multiple edge types allows the model to draw from these multiple relation types.

#### A.2.2 Node features

Each gene is associated with a four-view multi-omic vector that aggregates evidence about how the gene is *altered* or *active* in sixteen different cancer types: KIRC (kidney renal clear cell carcinoma), BRCA (breast invasive carcinoma), READ (rectum adenocarcinoma), PRAD (prostate adenocarcinoma), STAD (stomach adenocarcinoma), HNSC (head and neck squamous cell carcinoma), LUAD (lung adenocarcinoma), THCA (thyroid carcinoma), BLCA (bladder urothelial carcinoma), ESCA (esophageal carcinoma), LIHC (liver hepatocellular carcinoma), UCEC (uterine corpus endometrial carcinoma), COAD (colon adenocarcinoma), LUSC (lung squamous cell carcinoma), CESC (cervical squamous cell carcinoma and endocervical adenocarcinoma), and KIRP (kidney renal papillary cell carcinoma). This representation allows the model to exploit both pan-cancer regularities and tissue-specific idiosyncrasies in a unified space. The following omics types are included:

- **Copy-number alteration (CNA).** Chromosomal instability can duplicate or delete large DNA segments. We encode the resulting log_2_ copy-ratio for each gene. Amplifications drive oncogenes via dosage; deletions can inactivate tumour suppressors; either type of alteration increases the gene’s potential therapeutic relevance by changing pathway dynamics and dependencies.
- **Gene expression.** TPM-normalised RNA-seq counts serve as a proxy for transcriptional activity along the canoncial DNA → mRNA → protein axis. High expression marks pathway engagement and potential vulnerability; zero or strongly tissue-specific expression highlights candidates for of potential on-target toxicity.
- **Mutation frequency.** A *non-synonymous* variant changes an amino-acid and can alter protein function. We supply the fraction of tumours (TCGA) carrying at least one non-synonymous hit in each gene. Recurrent hits point to cancer *drivers*; high frequency therefore raises the prior that a gene is causally important – and a possible drug target.
- **Methylation.** CpG methylation at a promoter recruits proteins that compact chromatin and block transcription – known as *epigenetic silencing*. The *β* value (where 0 = unmethylated, 1 = fully methylated) distinguishes permanently “switched-off” genes from merely low-copy ones, helping the model avoid nominating silent targets.

##### Rationale and complementarity

Taken together, these four views cover structural (CNA), regulatory (expression, methylation), and genetic (mutation) evidence. This complementarity provides orthogonal signals that no single modality alone can provide, and enables the encoder to disambiguate mechanisms (e.g., high expression due to amplification versus loss of expression due to promoter hypermethylation).

##### Data source and reproducibility

We derive these features from TCGA, a widely used and rigorously curated resource for cancer genomics [35]. Its breadth, depth, and transparent processing pipelines enable reproducible comparisons across studies and typically provide stronger statistical power than smaller proprietary cohorts. While under-representation of rare histologies and understudied genes remains a limitation of any centralized resource, TCGA’s standardization and multi-omic scope make it an appropriate foundation for building generalizable target representations at scale.

##### Extensibility to additional modalities

MORGaN is feature-agnostic: any per-gene descriptor can be appended to the node feature vector without architectural changes. In particular, structural and chemoinformatics descriptors – such as binding-site fingerprints, pocket hydrophobicity, or docking-derived scores – are natural complements to biological priors. Embedding these signals would involve augmenting the node features with quantities derived from 3D structures or *in silico* screening. Because the present work focuses on *upstream* target prioritization from multi-omic and network context, a full end-to-end fusion with chemoinformatics is left for future work; we view this as an exciting extension toward unifying biological and chemical modalities in a single graph-learning pipeline.

#### A.2.3 Labels

##### Positive–unlabeled formulation

We frame the task as positive–unlabeled (PU) learning. High-confidence positives – FDA-approved or clinically validated drug targets – are known. However, *true* negatives do not exist: a gene without clinical evidence is not necessarily undruggable. To reflect this epistemic asymmetry, we treat the remaining genes as unlabeled and, for each train/validation/test split, sample negatives uniformly at random from this pool. This approach (i) avoids penalizing understudied genes, (ii) allows estimation of class-conditional risk without inventing a questionable negative set, and (iii) yields conservative evaluations because improvements must persist across independent negative samplings.

Moreover, drug target labels are intrinsically skewed (on the order of ~150 Tier-1 positives versus ~16,000 unlabeled genes). There is no authoritative set of genes that are *provably* undruggable, and previously intractable targets continue to become amenable with new modalities (e.g., PROTACs, molecular glues, mRNA therapeutics). We therefore create negatives by resampling a subset of unlabeled genes for every split:

- **Bias dilution.** Because the negative pool changes with each split, the classifier cannot overfit to idiosyncrasies of any single hand-curated list. Despite resampling, metric standard deviations remain low, indicating stable performance.
- **Graph neutrality.** Resampled negatives retain their full connectivity and multi-omic features, preserving the structural context established during pre-training. The model continues to learn from each gene’s neighbourhood and attributes even when a given gene is temporarily treated as negative, thereby avoiding topological artefacts that would arise from pruning or rewiring nodes.
- **Forward compatibility.** If a gene is later reclassified as drug target (e.g., due to a new modality), past experiments remain valid because that gene was never canonically fixed as negative. Benchmarks can be rerun with an updated label file without invalidating prior protocols.

These design choices mitigate pathway memorization, manage extreme class imbalance, and keep the evaluation protocol adaptable to methodological and pharmacological advances.

##### Why binary labels in practice

In principle, “druggability” spans a continuum of chemical tractability that evolves with technology. In practice, however, industrial target pipelines employ *discrete gates* (e.g., evidence of a small-molecule binder, a clinical candidate, or regulatory approval). We therefore label Tier-1 targets (approved or clinical candidates) as positives and sample negatives uniformly from unlabeled genes, mirroring how pipelines prioritize targets. This operational definition enables fair, reproducible benchmarking and aligns with prior work [4], while remaining compatible with future re-labeling as the field progresses.

## B Self-supervised masked pre-training

### Pre-training dynamics

Fig. 6 shows that the scaled-cosine reconstruction loss drops sharply during the first ten epochs, then converges smoothly, indicating that the model quickly captures first-order correlations and subsequently refines higher-order structure. The frozen embeddings obtained after 100 epochs serve as initialization for the downstream drug target classifier.

**Figure 6:**
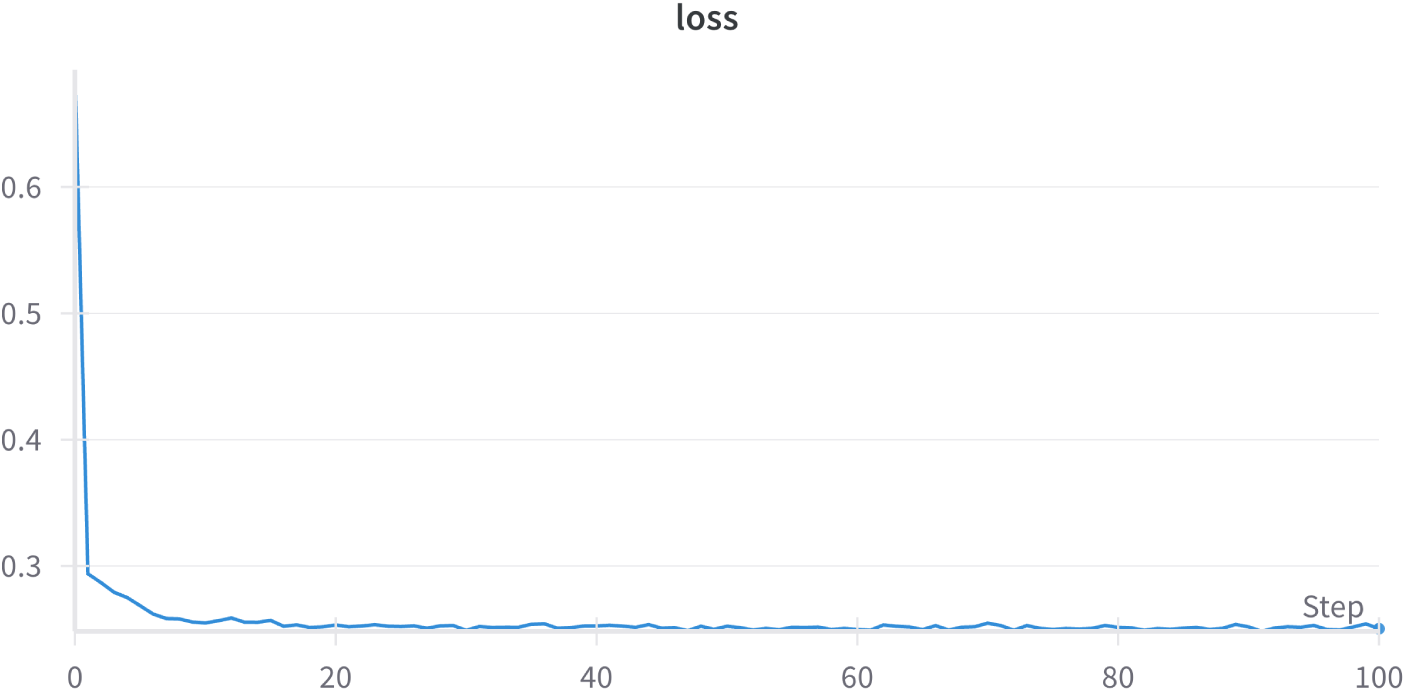
Scaled-cosine reconstruction loss during masked pre-training (mean ± s.d. over six splits).

### Hyper-parameter search in brief

A grid of 192 runs varied mask ratio (0.1–0.8), depth (1–4 RGCN layers), learning rate, weight decay and activation. The best AUPR clustered around a mask ratio of 0.5, two or three layers, PReLU activation, learning rate 10^−2^ for pre-training and 5×10^−3^ for fine-tuning, and weight decay 10^−3^ / 10^−4^ respectively (Fig. 7). These values constitute the default configuration shipped in the supplementary config.yaml; all reported results use that setting.

**Figure 7:**
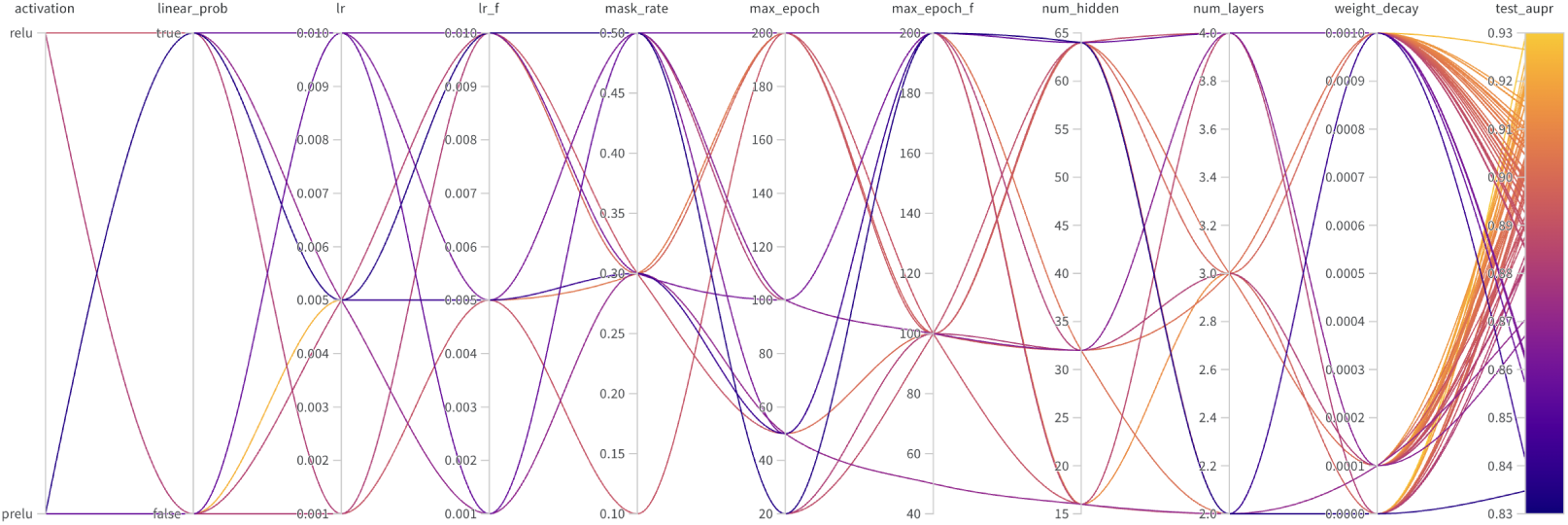
Parallel-coordinates view of the 192-run hyper-parameter sweep; colour encodes test AUPR. Orange lines highlight the high-performing region described in the text.

### Sensitivity to masking ratio

Masking ratio (feature corruption). Raising the fraction of masked features from 5% to 50% consistently improves downstream metrics, with AUPR rising by ∽ 4 pp and AUROC by ∽ 3 pp. A higher mask rate forces the encoder to rely more heavily on relational context instead of relying on raw features, leading to richer, more transferable embeddings. Beyond that, performance eventually degrades.

### Sensitivity to loss exponent

Increasing the loss exponent *γ* (the error curvature) in the SCE reconstruction loss steepens the penalty on large reconstruction errors. This gradually lifts AUPR from 0.907 (*γ* = 0.5) to 0.926 (*γ* = 5.0), but the gains are modest (*<* 2 pp AUPR over a ten-fold change) and all standard deviations overlap, indicating that the model remains broadly insensitive to the precise curvature of the loss.

Hence, performance improves with stronger feature corruption and a steeper loss, but the increments are small. Why such robustness? We believe that it can be traced back to two things:

1. **Aggregated objective.** The MAE sums residuals over six relation types and multidimensional features, so changing the weight on any individual error, via masking or *γ*, has a diluted global effect.
2. **Masking as a regulariser.** Even relatively moderate corruption (≥30%) regularises the model; once in this regime, additional changes are unlikely to reshape the learned space.

Practically, this means MORGaN can be deployed with default settings (e.g. 50% masking, *γ* = 3), still achieving within 1 − 2 pp of the best scores - greatly simplifying hyper-parameter tuning while underscoring the model’s inherent robustness.

### B.1 Defaults

- **Default hyperparameters:** mask ratio = 0.5, *γ*=3, 2–3 relation layers (PReLU), LR 1×10^−2^ (pre-train), 5×10^−3^ (fine-tune).
- **Early stopping:** monitor validation AUPR with patience 20 epochs.
- **Splits:** report mean ± s.d. over *k* seeds; use consistent positive fractions across splits.
- **Compute:** single spmm per layer via basis decomposition for efficiency; training-time wall clock improvements observed vs. per-relation updates.

## C Models

We keep the node features (§4.1) and the stratified 80/10/10 train–validation–test splits described in §4.4 the same across all models. Hyper-parameters are selected by grid search on the validation fold and seeds are fixed to 0, 1 for full reproducibility. Mean ± s.d. over six runs are reported in Table 3 of the main paper. The eight models fall into three tiers: feature-only, homogeneous (single-relation) graph and heterogeneous (multi-relation) graph.

### C.1 Feature-only models

#### Logistic regression

A linear classifier with *L*_2_ regularization trained on the node features, with no graph structure We use scikit-learn’s LogisticRegression(max_iter = 1000, penalty = “l2”, solver = “lbfgs”) and optimize the inverse regularization strength *C* over {0.01, 0.1, 1, 10}. Class weights are set inverse-frequency to counter the 1:1 positive/unlabeled sampling. This baseline tests whether a *strictly linear* decision boundary in feature space can already separate drug targets from non-drug targets.

#### Multilayer perceptron (MLP)

Identical input as above, but with two hidden layers to capture non-linear feature interactions. Architecture: [in → 64 → 32 → 2] with ReLU, dropout 0.2 after each hidden layer, and softmax output. Optimiser: Adam (lr = 1 × 10^−3^, weight-decay 5 × 10^−4^), batch size 256, 100 epochs, early stopping (patience 20). Validation tuning sweeps hidden size {32, 64, 128} and learning rate {1 × 10^−4^, 1 × 10^−3^, 5 × 10^−3^}. Serves as a capacity-matched non-graph baseline.

### C.2 Homogeneous-graph models

#### Graph convolutional network (GCN)

The vanilla spectral GCN operating on the *single* PPI edge set. Best configuration from the grid: two layers, hidden 128, PReLU activation, dropout 0.2, weight-decay 1 × 10^−4^. Input is a graph where nodes represent genes, node features are the same as above, and edges are derived from PPIs.

#### Graph attention network (GAT)

Multi-head attention on the same PPI graph. We use three layers with hidden 64 per head, LeakyReLU(0.2), feat-drop 0.2 and attn-drop 0.2. Heads are concatenated inside the network and averaged in the output layer. Edge-specific attention weights let the model down-weight noisy PPI links, providing a stronger yet still homogeneous comparator.

#### SMG-based (self-supervised masking)

Following Cui et al. [14] we add a masked-feature reconstruction pre-text stage to the GCN and GAT backbones. Mask ratio 0.5, 100 pre-training epochs (lr 1 × 10^−2^, weight-decay 1 × 10^−3^, cosine decay), then fine-tune as above for at most 200 epochs (lr 5 × 10^−3^). This pair isolates the effect of *self-supervision* while holding the single-relation topology constant.

### C.3 Heterogeneous-graph models

#### MODIG

The multi-omics, multi-relation GAT of Zhao et al. [21] trained on our six-edge-type graph. Each relation is processed by its own two-layer GAT; relation-specific embeddings are fused with learned view-level attention before a final MLP classifier. We keep the authors’ recommended settings (hidden 128, 8 heads, dropout 0.3) and tune only the learning rate. MODIG gauges the benefit of heterogeneous edges without any self-supervised pre-training.

#### MDMNI-DGD

The meta-path DNN of Li et al. [22] – a six-view extension of MODIG that stacks dense layers on hand-crafted meta-path incidence vectors. We train the model on our dataset, following the original paper – we use three hidden layers (256–128–64, dropout 0.3) and Adam (lr 1 × 10^−3^). This baseline retains heterogeneous information but replaces GNN message passing with fully-connected fusion, testing whether explicit relational reasoning is needed.

Together, these baselines allus us to disentangle the contributions of (i) multi-omic feature depth, (ii) homogeneous versus heterogeneous topology, and (iii) self-supervised pre-training, ultimately demonstrating the incremental value added by each MORGaN component.

## D Computational requirements and efficiency

All timings were obtained on a **MacBook Pro (Apple M3, 8-core CPU, 16 GB RAM, macOS 15.4.1)** with no GPU acceleration. Table 2 compares MORGaN to the two strongest heterogeneous baselines.

**Table 2:** Runtime on the six-relation graph (mean ± s.d. over six runs).

| Model | CPU time / epoch (s) | End-to-end time (s) |
| --- | --- | --- |
| MODIG | 18.60 ± 1.18 | 1 582 ± 116 |
| MDMNI-DGD | 5.69 ± 0.82 | 566 ± 16 |
| **MORGaN** | **0.23 ± 0.07** | **24.3 ± 2.9** |

**Key numbers.** MORGaN trains ≈80× **faster per epoch** than MODIG and completes the full pre-train + fine-tune pipeline ≈65× **faster**. Put differently, a hyper-parameter sweep that takes one day with MODIG finishes in under 30 minutes with MORGaN on a traditional laptop.

## E Results table

**Table 3:** Test-set performance of MORGaN versus eight models on the drug target identification task (mean ± s.d.). **Bold** numbers indicate the best score per column; *italic* numbers mark the second best. All models receive the same multi-omic node features; heterogeneous methods (bottom blocks) also share the identical six-relation graph. Same splits. A full description of the baselines is provided in Appendix C.

| Model | AUPR | AUROC | Accuracy | F1 Score |
| --- | --- | --- | --- | --- |
| Logistic regression | 0.749 ± 0.045 | 0.682 ± 0.055 | 0.620 ± 0.044 | 0.577 ± 0.096 |
| MLP | 0.675 ± 0.052 | 0.722 ± 0.045 | 0.722 ± 0.045 | 0.701 ± 0.035 |
| GCN | 0.721 ± 0.020 | 0.766 ± 0.005 | 0.715 ± 0.025 | 0.722 ± 0.037 |
| GAT | 0.699 ± 0.005 | 0.764 ± 0.009 | 0.724 ± 0.015 | 0.742 ± 0.005 |
| SMG-GCN [14] | 0.714 ± 0.009 | 0.763 ± 0.011 | 0.729 ± 0.029 | 0.724 ± 0.014 |
| SMG-GAT [14] | 0.708 ± 0.005 | 0.776 ± 0.005 | 0.732 ± 0.027 | 0.751 ± 0.014 |
| MODIG [21] | 0.764 ± 0.017 | 0.837 ± 0.015 | 0.794 ± 0.009 | 0.810 ± 0.007 |
| MDMNI-DGD [22] | 0.815 ± 0.019 | 0.877 ± 0.003 | 0.664 ± 0.038 | 0.741 ± 0.022 |
| MORGaN (no pre-training) | *0.879 ± 0.006* | *0.900 ± 0.006* | *0.898 ± 0.006* | *0.902 ± 0.006* |
| **MORGaN (with pre-training)** | **0.888 ± 0.004** | **0.907 ± 0.005** | **0.915 ± 0.005** | **0.917 ± 0.004** |

## F Ablation studies

The main paper shows that MORGaN outperforms eight strong baselines; the natural follow-up question is *why*. We therefore conduct six systematic ablation experiments, which all run on the same train–validation–test splits and are evaluated with the same metrics as the main results (AUPR is the headline score).

1. **PPI-source comparisons** (Table 4) swap the base PPI layer among five popular databases (STRING-db, PCNet, CPDB, IRefIndex 2015, IRefIndex 2009) while holding all other relations and features constant. CPDB is used in all main-paper experiments.
2. **Feature ablations** (Table 5) isolate the importance of the four node-feature modalities (CNA, gene expression, methylation, mutation frequency) by training MORGaN on every single, pairwise, triple, and full combination.
3. **Edge-type ablations** (Tables 6-7) repeat the experiment for the six biological relation types.
4. **Randomized-edge control ablations** (Tables 8-9) replace each real edge set with a degree-preserved shuffle keeping node features unchanged. Performance dropping to chance under this perturbation demonstrates that the improvements arise from genuine biology rather than increased edge density or model capacity.
5. **Domain-restricted (organ-system) training** (Table 10) tests whether pan-cancer gains arise from cross-tumour transfer or from a few dominant entities. We retrain MORGaN on *organ-specific feature and label subsets* while holding graph topology fixed.
6. **Model ablations.** We swap the basis-decomposed RGCN encoder for a relational GIN (RGIN) with matched depth/width/parameters and identical pre-training task, decoder, and schedule to probe whether gains are operator-specific or persist across encoder families (Table 11). In addition, we ablate the efficiency components – vertical stacking and weight decomposition – showing that stacking provides the dominant speedup while decomposition preserves this throughput, reduces parameters via sharing, and acts as a mild regularizer (Table 12).

All ablation results are averaged over the six stratified shuffle–split runs described in §4.4; one standard deviation is shown for completeness. The next subsections present the detailed numbers and summarize the key observations.

### F.1 Comparison between PPI datasets

**Table 4:** PPI-source comparison. Performance of MORGaN when the PPI layer is sourced from five popular interaction databases. All other edge types and node features are kept identical. **Bold** numbers highlight the best score *within each column*. STRING-db provides the most informative PPI set, pushing AUPR to 0.971, whereas the older IRefIndex releases yield lower accuracy despite comparable AUPR/AUROC figures.

| Features | AUPR | AUROC | Accuracy | F1 |
| --- | --- | --- | --- | --- |
| CPDB | 0.888 ± 0.004 | 0.906 ± 0.004 | 0.917 ± 0.004 | 0.919 ± 0.004 |
| IRefIndex 2015 | 0.949 ± 0.008 | 0.944 ± 0.004 | 0.866 ± 0.011 | 0.869 ± 0.010 |
| IRefIndex | 0.949 ± 0.008 | 0.944 ± 0.004 | 0.866 ± 0.011 | 0.869 ± 0.010 |
| PCNet | 0.950 ± 0.008 | 0.941 ± 0.007 | 0.893 ± 0.004 | 0.888 ± 0.004 |
| STRINGdb | **0.971 ± 0.002** | **0.970 ± 0.001** | **0.927 ± 0.007** | **0.927 ± 0.007** |

### F.2 Feature ablations

**Table 5:** Ablation of the four input omics modalities. . Blocks separated by lines correspond to (top to bottom) single-, pair-, triple- and four-modality configurations. **Bold** numbers highlight the best score within each column, and *italics* highlight the second-best. Copy-number alterations (CNA) are the most informative modality on their own, whereas combining CNA with gene expression (GE) or mutation frequency (MF) restores accuracy and F1 to the highest levels. Using all four modalities yields a balanced performance but does not surpass the best CNA–based subsets on AUPR.

| Features | AUPR | AUROC | Accuracy | F1 |
| --- | --- | --- | --- | --- |
| Copy Number Alterations (CNA) | **0.908 ± 0.002** | *0.927 ± 0.001* | 0.907 ± 0.005 | 0.909 ± 0.005 |
| Gene Expression (GE) | 0.859 ± 0.007 | 0.919 ± 0.002 | 0.917 ± 0.004 | 0.919 ± 0.004 |
| Methylation (METH) | 0.884 ± 0.003 | 0.907 ± 0.002 | 0.913 ± 0.004 | 0.914 ± 0.004 |
| Mutation Frequency (MF) | 0.866 ± 0.002 | 0.909 ± 0.002 | 0.900 ± 0.008 | 0.902 ± 0.007 |
| CNA + GE | 0.874 ± 0.011 | 0.920 ± 0.007 | **0.919 ± 0.000** | **0.921 ± 0.000** |
| CNA + METH | 0.893 ± 0.005 | 0.910 ± 0.007 | 0.909 ± 0.004 | 0.911 ± 0.004 |
| CNA + MF | **0.908 ± 0.004** | **0.929 ± 0.003** | 0.911 ± 0.000 | 0.913 ± 0.000 |
| GE + METH | 0.891 ± 0.018 | 0.907 ± 0.002 | 0.909 ± 0.008 | 0.911 ± 0.007 |
| GE + MF | 0.881 ± 0.005 | 0.920 ± 0.002 | 0.917 ± 0.004 | 0.918 ± 0.004 |
| METH + MF | 0.890 ± 0.004 | 0.909 ± 0.001 | 0.915 ± 0.005 | 0.917 ± 0.005 |
| CNA + GE + METH | 0.891 ± 0.002 | 0.912 ± 0.002 | 0.917 ± 0.004 | 0.919 ± 0.004 |
| CNA + GE + MF | 0.886 ± 0.006 | 0.918 ± 0.004 | **0.919 ± 0.000** | **0.921 ± 0.000** |
| CNA + METH + MF | 0.897 ± 0.005 | 0.916 ± 0.006 | 0.911 ± 0.007 | 0.913 ± 0.006 |
| GE + METH + MF | **0.908 ± 0.006** | 0.913 ± 0.007 | 0.896 ± 0.008 | 0.901 ± 0.007 |
| CNA + GE + METH + MF | 0.888 ± 0.005 | 0.906 ± 0.004 | 0.917 ± 0.004 | 0.919 ± 0.004 |

### F.3 Edge ablations

**Table 6:** Edge–type ablation. , part I (up to four relation types). Each row shows test performance when the heterogeneous graph is restricted to the specified subset of biological relations. Values are mean ± s.d. over the six splits described in §4.4. The full six-relation result (AUPR = 0.888, cf. Table 3) is given for reference in Table 7. The horizontal rules separate 1-, 2-, 3- and 4-relation configurations. **Bold** numbers mark the best score within each block.

| Relations | AUPR | AUROC | Accuracy | F1 |
| --- | --- | --- | --- | --- |
| Co-expression (Coexpr.) | 0.788 ± 0.006 | 0.788 ± 0.003 | 0.805 ± 0.000 | 0.782 ± 0.000 |
| Domain Similarity (DomSim) | 0.764 ± 0.000 | 0.533 ± 0.000 | 0.537 ± 0.000 | 0.123 ± 0.000 |
| GO Semantic Similarity (GO) | **0.883 ± 0.002** | 0.819 ± 0.001 | 0.805 ± 0.000 | 0.778 ± 0.000 |
| Pathway Co-occurrence (Path) | 0.863 ± 0.008 | **0.847 ± 0.004** | **0.843 ± 0.004** | **0.832 ± 0.005** |
| Sequence Similarity (SeqSim) | 0.752 ± 0.000 | 0.508 ± 0.000 | 0.512 ± 0.000 | 0.032 ± 0.000 |
| Coexpr. + DomSim | 0.809 ± 0.007 | 0.798 ± 0.002 | 0.813 ± 0.000 | 0.793 ± 0.000 |
| Coexpr. + GO | 0.882 ± 0.015 | 0.892 ± 0.003 | 0.878 ± 0.000 | 0.878 ± 0.001 |
| Coexpr. + PPI | 0.827 ± 0.021 | 0.881 ± 0.001 | 0.835 ± 0.015 | 0.825 ± 0.013 |
| Coexpr. + Path. | 0.878 ± 0.007 | **0.904 ± 0.003** | **0.894 ± 0.011** | **0.894 ± 0.013** |
| Coexpr. + SeqSim | 0.783 ± 0.001 | 0.790 ± 0.001 | 0.805 ± 0.000 | 0.782 ± 0.000 |
| DomSim + GO | 0.887 ± 0.008 | 0.825 ± 0.015 | 0.805 ± 0.000 | 0.778 ± 0.000 |
| DomSim + PPI | 0.756 ± 0.007 | 0.804 ± 0.004 | 0.738 ± 0.041 | 0.740 ± 0.068 |
| DomSim + Path. | 0.878 ± 0.003 | 0.855 ± 0.007 | 0.841 ± 0.008 | 0.830 ± 0.010 |
| DomSim + SeqSim | 0.769 ± 0.000 | 0.541 ± 0.000 | 0.545 ± 0.000 | 0.152 ± 0.000 |
| GO + PPI | 0.849 ± 0.030 | 0.789 ± 0.074 | 0.813 ± 0.016 | 0.792 ± 0.028 |
| GO + Path. | **0.904 ± 0.003** | 0.901 ± 0.001 | **0.894 ± 0.000** | **0.894 ± 0.001** |
| GO + SeqSim | 0.880 ± 0.004 | 0.812 ± 0.002 | 0.805 ± 0.000 | 0.778 ± 0.000 |
| PPI + Path. | 0.869 ± 0.007 | **0.904 ± 0.002** | 0.843 ± 0.004 | 0.835 ± 0.006 |
| PPI + SeqSim | 0.740 ± 0.014 | 0.789 ± 0.014 | 0.726 ± 0.043 | 0.729 ± 0.059 |
| Path. + SeqSim | 0.853 ± 0.004 | 0.839 ± 0.008 | 0.835 ± 0.004 | 0.822 ± 0.005 |
| Coexpr. + DomSim + GO | 0.897 ± 0.010 | 0.895 ± 0.002 | 0.878 ± 0.007 | 0.877 ± 0.008 |
| Coexpr. + DomSim + PPI | 0.856 ± 0.015 | 0.895 ± 0.002 | 0.833 ± 0.019 | 0.826 ± 0.018 |
| Coexpr. + DomSim + Path. | 0.861 ± 0.023 | 0.902 ± 0.004 | 0.909 ± 0.004 | 0.910 ± 0.004 |
| Coexpr. + DomSim + SeqSim | 0.811 ± 0.004 | 0.802 ± 0.003 | 0.813 ± 0.000 | 0.793 ± 0.000 |
| Coexpr. + GO + PPI | 0.903 ± 0.018 | 0.909 ± 0.033 | 0.876 ± 0.004 | 0.876 ± 0.005 |
| Coexpr. + GO + Path. | 0.891 ± 0.001 | **0.918 ± 0.001** | **0.917 ± 0.004** | **0.918 ± 0.004** |
| Coexpr. + GO + SeqSim | 0.890 ± 0.025 | 0.881 ± 0.009 | 0.872 ± 0.004 | 0.870 ± 0.004 |
| Coexpr. + PPI + Path. | 0.878 ± 0.005 | 0.917 ± 0.003 | 0.909 ± 0.004 | 0.910 ± 0.004 |
| Coexpr. + PPI + SeqSim | 0.827 ± 0.015 | 0.874 ± 0.004 | 0.819 ± 0.010 | 0.810 ± 0.021 |
| Coexpr. + Path. + SeqSim | 0.866 ± 0.032 | 0.900 ± 0.003 | 0.909 ± 0.004 | 0.910 ± 0.004 |
| DomSim + GO + PPI | 0.861 ± 0.031 | 0.800 ± 0.068 | 0.807 ± 0.004 | 0.782 ± 0.008 |
| DomSim + GO + Path. | **0.910 ± 0.001** | 0.910 ± 0.000 | 0.902 ± 0.000 | 0.903 ± 0.000 |
| DomSim + GO + SeqSim | 0.892 ± 0.004 | 0.835 ± 0.009 | 0.811 ± 0.004 | 0.786 ± 0.006 |
| DomSim + PPI + Path. | 0.886 ± 0.007 | 0.914 ± 0.008 | 0.860 ± 0.012 | 0.852 ± 0.015 |
| DomSim + PPI + SeqSim | 0.778 ± 0.006 | 0.810 ± 0.006 | 0.754 ± 0.008 | 0.766 ± 0.009 |
| DomSim + Path. + SeqSim | 0.875 ± 0.005 | 0.856 ± 0.007 | 0.848 ± 0.008 | 0.837 ± 0.010 |
| GO + PPI + Path. | 0.895 ± 0.005 | 0.889 ± 0.014 | 0.894 ± 0.000 | 0.894 ± 0.000 |
| GO + PPI + SeqSim | 0.872 ± 0.029 | 0.850 ± 0.066 | 0.797 ± 0.022 | 0.779 ± 0.007 |
| GO + Path. + SeqSim | 0.902 ± 0.002 | 0.901 ± 0.001 | 0.894 ± 0.000 | 0.894 ± 0.000 |
| PPI + Path. + SeqSim | 0.878 ± 0.009 | 0.905 ± 0.007 | 0.850 ± 0.008 | 0.840 ± 0.010 |
| Coexpr. + DomSim + GO + PPI | 0.905 ± 0.018 | 0.907 ± 0.031 | 0.882 ± 0.008 | 0.882 ± 0.007 |
| Coexpr. + DomSim + GO + Path | 0.899 ± 0.001 | **0.920 ± 0.001** | **0.919 ± 0.000** | **0.921 ± 0.000** |
| Coexpr. + DomSim + GO + SeqSim | 0.898 ± 0.010 | 0.878 ± 0.013 | 0.870 ± 0.015 | 0.868 ± 0.016 |
| Coexpr. + DomSim + PPI + Path. | 0.870 ± 0.030 | 0.905 ± 0.016 | 0.909 ± 0.004 | 0.910 ± 0.004 |
| Coexpr. + DomSim + PPI + SeqSim | 0.844 ± 0.026 | 0.885 ± 0.010 | 0.833 ± 0.010 | 0.837 ± 0.007 |
| Coexpr. + DomSim + Path. + SeqSim | 0.863 ± 0.022 | 0.902 ± 0.005 | 0.909 ± 0.004 | 0.910 ± 0.004 |
| Coexpr. + GO + PPI + Path. | 0.896 ± 0.003 | 0.915 ± 0.003 | 0.915 ± 0.008 | 0.917 ± 0.008 |
| Coexpr. + GO + PPI + SeqSim | 0.876 ± 0.002 | 0.861 ± 0.001 | 0.880 ± 0.010 | 0.879 ± 0.010 |
| Coexpr. + GO + Path. + SeqSim | 0.884 ± 0.003 | 0.915 ± 0.000 | **0.919 ± 0.000** | **0.921 ± 0.000** |
| Coexpr. + PPI + Path. + SeqSim | 0.866 ± 0.022 | 0.900 ± 0.004 | 0.904 ± 0.012 | 0.906 ± 0.012 |

**Table 7:**
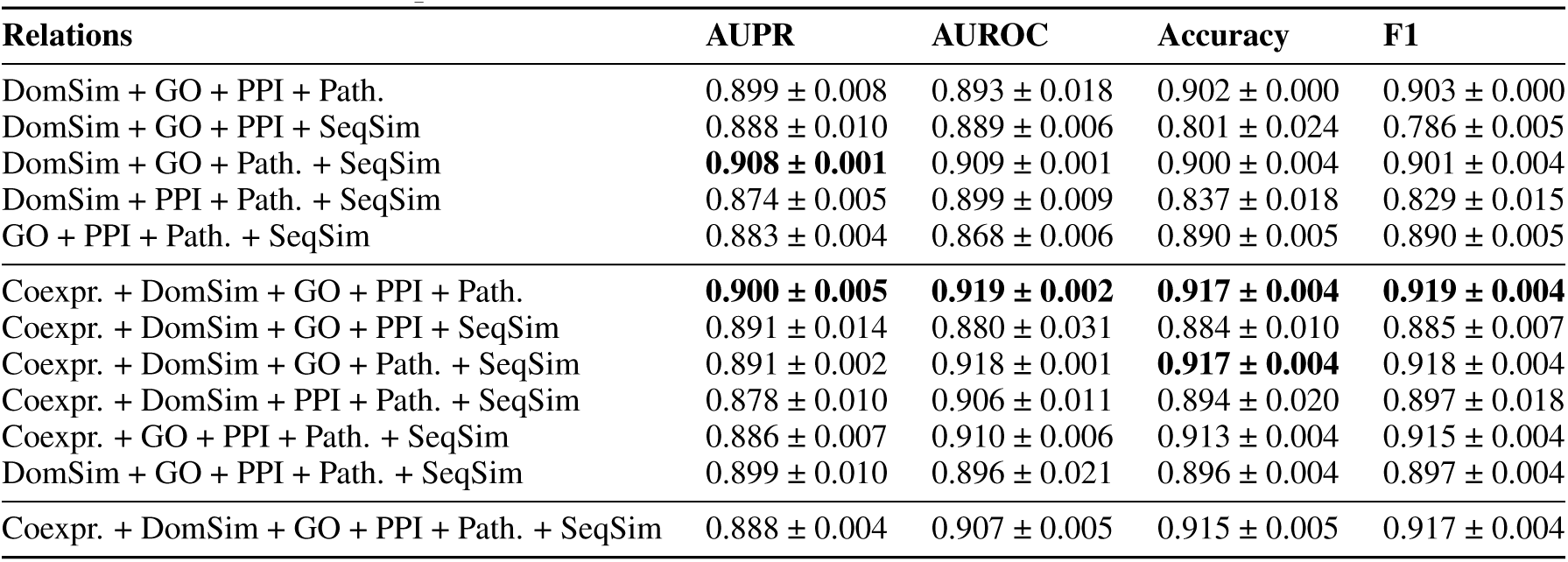
Edge–type ablation. , part II (four to six relation types, continued). This table completes the sweep by listing the remaining four- and five-relation subsets followed by the full six-relation graph (bottom row). Metrics are reported as mean ± s.d. over six runs.

### F.4 Edge ablations (randomized)

**Table 8:** Randomized–edge ablation. , part I (up to three relation types). For each subset of biological relations we replace every edge with a degree-preserved shuffle, keeping node features unchanged. Performance collapses to chance level (AUPR ≈ 0.5, AUROC ≈ 0.5), demonstrating that MORGaN’s gains in Table 6 come from *biologically meaningful* topology rather than edge density or parameter count. Horizontal rules separate 1-, 2- and 3-relation configurations; values are mean ± s.d. over six runs.

| Relations | AUPR | AUROC | Accuracy | F1 |
| --- | --- | --- | --- | --- |
| Coexpr. | 0.512 ± 0.050 | 0.499 ± 0.060 | 0.503 ± 0.046 | 0.494 ± 0.072 |
| DomSim | 0.372 ± 0.141 | 0.496 ± 0.005 | 0.504 ± 0.000 | 0.349 ± 0.367 |
| GO | 0.548 ± 0.036 | 0.531 ± 0.039 | 0.520 ± 0.045 | 0.482 ± 0.112 |
| Path. | 0.530 ± 0.038 | 0.500 ± 0.018 | 0.502 ± 0.015 | 0.479 ± 0.157 |
| SeqSim | 0.498 ± 0.289 | 0.496 ± 0.005 | 0.498 ± 0.004 | 0.166 ± 0.332 |
| Coexpr. + DomSim | 0.480 ± 0.040 | 0.494 ± 0.053 | 0.488 ± 0.018 | 0.491 ± 0.059 |
| Coexpr. + GO | 0.504 ± 0.026 | 0.514 ± 0.022 | 0.528 ± 0.018 | 0.504 ± 0.078 |
| Coexpr. + PPI | 0.622 ± 0.148 | 0.494 ± 0.025 | 0.504 ± 0.007 | 0.141 ± 0.164 |
| Coexpr. + Path. | 0.583 ± 0.030 | 0.579 ± 0.019 | 0.553 ± 0.015 | 0.575 ± 0.027 |
| Coexpr. + SeqSim | 0.469 ± 0.044 | 0.460 ± 0.071 | 0.480 ± 0.043 | 0.482 ± 0.060 |
| DomSim + GO | 0.538 ± 0.054 | 0.515 ± 0.040 | 0.520 ± 0.015 | 0.494 ± 0.151 |
| DomSim + PPI | 0.582 ± 0.119 | 0.520 ± 0.047 | 0.512 ± 0.033 | 0.396 ± 0.287 |
| DomSim + Path. | 0.491 ± 0.013 | 0.473 ± 0.018 | 0.492 ± 0.005 | 0.341 ± 0.168 |
| DomSim + SeqSim | 0.628 ± 0.145 | 0.512 ± 0.014 | 0.508 ± 0.014 | 0.669 ± 0.006 |
| GO + PPI | 0.619 ± 0.149 | 0.495 ± 0.014 | 0.508 ± 0.008 | 0.166 ± 0.261 |
| GO + Path. | 0.503 ± 0.042 | 0.474 ± 0.052 | 0.484 ± 0.024 | 0.478 ± 0.094 |
| GO + SeqSim | 0.483 ± 0.042 | 0.476 ± 0.027 | 0.496 ± 0.034 | 0.358 ± 0.159 |
| PPI + Path. | 0.640 ± 0.134 | 0.497 ± 0.047 | 0.502 ± 0.040 | 0.359 ± 0.311 |
| PPI + SeqSim | 0.634 ± 0.132 | 0.525 ± 0.040 | 0.512 ± 0.022 | 0.337 ± 0.371 |
| Path. + SeqSim | 0.488 ± 0.025 | 0.485 ± 0.037 | 0.494 ± 0.023 | 0.473 ± 0.172 |
| Coexpr. + DomSim + GO | 0.539 ± 0.023 | 0.524 ± 0.032 | 0.514 ± 0.012 | 0.496 ± 0.026 |
| Coexpr. + DomSim + PPI | 0.492 ± 0.012 | 0.487 ± 0.009 | 0.498 ± 0.004 | 0.479 ± 0.071 |
| Coexpr. + DomSim + Path. | 0.507 ± 0.018 | 0.501 ± 0.028 | 0.520 ± 0.015 | 0.519 ± 0.023 |
| Coexpr. + DomSim + SeqSim | 0.494 ± 0.015 | 0.478 ± 0.034 | 0.482 ± 0.032 | 0.463 ± 0.078 |
| Coexpr. + GO + PPI | 0.617 ± 0.066 | 0.590 ± 0.067 | 0.549 ± 0.062 | 0.561 ± 0.053 |
| Coexpr. + GO + Path. | 0.465 ± 0.045 | 0.463 ± 0.082 | 0.480 ± 0.072 | 0.438 ± 0.081 |
| Coexpr. + GO + SeqSim | 0.566 ± 0.048 | 0.549 ± 0.055 | 0.551 ± 0.033 | 0.564 ± 0.030 |
| Coexpr. + PPI + Path. | 0.480 ± 0.022 | 0.485 ± 0.031 | 0.472 ± 0.027 | 0.468 ± 0.177 |
| Coexpr. + PPI + SeqSim | 0.465 ± 0.030 | 0.447 ± 0.051 | 0.467 ± 0.060 | 0.395 ± 0.113 |
| Coexpr. + Path. + SeqSim | 0.487 ± 0.036 | 0.493 ± 0.037 | 0.504 ± 0.016 | 0.541 ± 0.087 |

**Table 9:** Randomized–edge ablation. , part II (three to six relation types). Continuation of Table 8, covering the remaining three-, four-, five- and full six-relation shuffles. Even with all six relation layers present but randomized, MORGaN remains close to random guessing, reinforcing that the real multi-relation structure (Table 7) is essential for predictive power.

| Relations | AUPR | AUROC | Accuracy | F1 |
| --- | --- | --- | --- | --- |
| DomSim + GO + PPI | 0.503 ± 0.051 | 0.522 ± 0.061 | 0.490 ± 0.029 | 0.467 ± 0.152 |
| DomSim + GO + Path. | 0.503 ± 0.048 | 0.531 ± 0.057 | 0.537 ± 0.040 | 0.546 ± 0.079 |
| DomSim + GO + SeqSim | 0.532 ± 0.026 | 0.509 ± 0.020 | 0.498 ± 0.017 | 0.552 ± 0.115 |
| DomSim + PPI + Path. | 0.582 ± 0.131 | 0.518 ± 0.063 | 0.510 ± 0.028 | 0.420 ± 0.316 |
| DomSim + PPI + SeqSim | 0.532 ± 0.008 | 0.537 ± 0.022 | 0.533 ± 0.041 | 0.432 ± 0.224 |
| DomSim + Path. + SeqSim | 0.498 ± 0.050 | 0.523 ± 0.047 | 0.524 ± 0.037 | 0.489 ± 0.132 |
| GO + PPI + Path. | 0.491 ± 0.063 | 0.470 ± 0.075 | 0.480 ± 0.060 | 0.471 ± 0.176 |
| GO + PPI + SeqSim | 0.506 ± 0.049 | 0.481 ± 0.034 | 0.486 ± 0.026 | 0.472 ± 0.114 |
| GO + Path. + SeqSim | 0.504 ± 0.028 | 0.524 ± 0.052 | 0.533 ± 0.030 | 0.535 ± 0.074 |
| PPI + Path. + SeqSim | 0.552 ± 0.040 | 0.555 ± 0.023 | 0.539 ± 0.046 | 0.504 ± 0.127 |
| Coexpr. + DomSim + GO + PPI | 0.529 ± 0.070 | 0.494 ± 0.064 | 0.504 ± 0.083 | 0.522 ± 0.079 |
| Coexpr. + DomSim + GO + Path. | 0.532 ± 0.079 | 0.549 ± 0.102 | 0.541 ± 0.060 | 0.550 ± 0.054 |
| Coexpr. + DomSim + GO + SeqSim | 0.486 ± 0.045 | 0.489 ± 0.046 | 0.502 ± 0.031 | 0.465 ± 0.081 |
| Coexpr. + DomSim + PPI + Path. | 0.549 ± 0.053 | 0.551 ± 0.019 | 0.549 ± 0.037 | 0.460 ± 0.288 |
| Coexpr. + DomSim + PPI + SeqSim | 0.477 ± 0.018 | 0.443 ± 0.023 | 0.470 ± 0.022 | 0.488 ± 0.163 |
| Coexpr. + DomSim + Path. + SeqSim | 0.520 ± 0.053 | 0.461 ± 0.029 | 0.490 ± 0.014 | 0.449 ± 0.041 |
| Coexpr. + GO + PPI + Path. | 0.526 ± 0.021 | 0.504 ± 0.047 | 0.496 ± 0.026 | 0.271 ± 0.076 |
| Coexpr. + GO + PPI + SeqSim | 0.496 ± 0.081 | 0.475 ± 0.087 | 0.470 ± 0.027 | 0.509 ± 0.227 |
| Coexpr. + GO + Path. + SeqSim | 0.516 ± 0.040 | 0.507 ± 0.048 | 0.520 ± 0.030 | 0.486 ± 0.049 |
| Coexpr. + PPI + Path. + SeqSim | 0.538 ± 0.036 | 0.554 ± 0.026 | 0.541 ± 0.017 | 0.467 ± 0.172 |
| DomSim + GO + PPI + Path. | 0.519 ± 0.050 | 0.507 ± 0.040 | 0.512 ± 0.037 | 0.539 ± 0.094 |
| DomSim + GO + PPI + SeqSim | 0.529 ± 0.039 | 0.531 ± 0.026 | 0.533 ± 0.014 | 0.524 ± 0.093 |
| DomSim + GO + Path. + SeqSim | 0.585 ± 0.067 | 0.582 ± 0.061 | 0.561 ± 0.049 | 0.547 ± 0.037 |
| DomSim + PPI + Path. + SeqSim | 0.537 ± 0.066 | 0.536 ± 0.057 | 0.518 ± 0.017 | 0.524 ± 0.086 |
| GO + PPI + Path. + SeqSim | 0.481 ± 0.049 | 0.468 ± 0.055 | 0.492 ± 0.045 | 0.462 ± 0.151 |
| Coexpr. + DomSim + GO + PPI + Path. | 0.506 ± 0.019 | 0.536 ± 0.014 | 0.520 ± 0.040 | 0.493 ± 0.236 |
| Coexpr. + DomSim + GO + PPI + SeqSim | 0.511 ± 0.048 | 0.507 ± 0.038 | 0.496 ± 0.000 | 0.463 ± 0.095 |
| Coexpr. + DomSim + GO + Path. + SeqSim | 0.503 ± 0.011 | 0.491 ± 0.011 | 0.504 ± 0.018 | 0.525 ± 0.087 |
| Coexpr. + DomSim + PPI + Path. + SeqSim | 0.460 ± 0.031 | 0.449 ± 0.039 | 0.472 ± 0.040 | 0.405 ± 0.092 |
| Coexpr. + GO + PPI + Path. + SeqSim | 0.515 ± 0.051 | 0.507 ± 0.052 | 0.506 ± 0.054 | 0.341 ± 0.262 |
| DomSim + GO + PPI + Path. + SeqSim | 0.477 ± 0.012 | 0.469 ± 0.016 | 0.484 ± 0.019 | 0.419 ± 0.221 |
| Coexpr. + DomSim + GO + PPI + Path. + SeqSim | 0.505 ± 0.022 | 0.479 ± 0.036 | 0.480 ± 0.038 | 0.472 ± 0.122 |

### F.5 Domain-restricted (organ-system) training

To determine whether MORGaN’s accuracy is driven by a handful of tumor entities or is truly pan-cancer, we trained six separate models, each restricted to one “organ-system” (omics features retained only for the cancer types listed in brackets), based on those already included in the pan-cancer feature set used to train our original model:

- Head and Neck [HNSC]
- Gastro-intestinal [ESCA, STAD, LIHC, COAD, READ]
- Respiratory [LUAD, LUSC]
- Genitourinary [KIRC, KIRP, BLCA, PRAD]
- Reproductive [UCEC, CESC, BRCA]
- Endocrine [THCA]

The table below reports mean ± s.d. over three random splits (70*/*15*/*15%).

Across the profiled organ systems, performance is uniformly strong (AUPR = 0.892 − 0.919; AUROC = 0.891 − 0.927; Acc = 0.874 − 0.905; F1 = 0.877 − 0.908), indicating that MORGaN’s accuracy is not driven by a single tissue context. Variation is modest (absolute AUPR spread ≤ 0.027 with s.d. ≤ 0.016), and tracks data availability: the Respiratory group achieves the highest scores (AUPR 0.919 ± 0.003, AUROC 0.927 ± 0.002), while Head & Neck and Reproductive, which have fewer established positives, are slightly lower but remain well within the high-performing regime (AUPR ≈ 0.892 − 0.893, AUROC ≈ 0.891 − 0.901). Gastrointestinal and Genitourinary are consistently competitive (e.g., AUPR 0.898 and 0.910; AUROC 0.913 and 0.922, respectively). In short, MORGaN generalizes across cancer types; although joint pan-cancer training yields the single best overall model, the per-tissue experiments show that it retains high fidelity even when feature sets are restricted to smaller, system-specific vectors.

**Table 10:** Performance by tissue group (mean ± s.d.).

| Tissue group | AUPR | AUROC | Accuracy | F1 |
| --- | --- | --- | --- | --- |
| Gastrointestinal | $0.898 \pm 0.003$ | $0.913 \pm 0.005$ | $0.892 \pm 0.004$ | $0.896 \pm 0.004$ |
| Respiratory | $0.919 \pm 0.003$ | $0.927 \pm 0.002$ | $0.890 \pm 0.004$ | $0.896 \pm 0.005$ |
| Head and neck | $0.893 \pm 0.008$ | $0.901 \pm 0.003$ | $0.874 \pm 0.012$ | $0.877 \pm 0.016$ |
| Genitourinary | $0.910 \pm 0.009$ | $0.922 \pm 0.010$ | $0.905 \pm 0.004$ | $0.908 \pm 0.005$ |
| Reproductive | $0.892 \pm 0.007$ | $0.891 \pm 0.007$ | $0.874 \pm 0.009$ | $0.877 \pm 0.010$ |

### F.6 Model ablations

#### F.6.1 Encoder-family

To test whether MORGaN’s gains depend on the specific relational operator, we replace the basis-decomposed **RGCN** encoder/decoder with a **Relational GIN (RGIN)** backbone while keeping the pre-training objective, decoder head, data splits, optimization schedule, and regularization unchanged. We match depth/width to keep parameter count and per-epoch compute comparable.

Table 11 reports mean ± s.d. across the same splits used elsewhere. RGIN performance is comparable with that achieved by our RGCN configuration, indicating that MORGaN’s gains primarily arise from the multi-relation masking objective and the information in the heterogeneous graph rather than from a particular choice of message-passing operator.

**Table 11:** Encoder-family ablation: replacing RGCN with RGIN inside MORGaN (mean ± s.d. across identical splits).

|  | AUPR | AUROC | Accuracy | F1 |
| --- | --- | --- | --- | --- |
| RGCN | $0.888 \pm 0.004$ | $0.907 \pm 0.005$ | $0.915 \pm 0.005$ | $0.917 \pm 0.004$ |
| RGIN | $0.908 \pm 0.005$ | $0.913 \pm 0.011$ | $0.898 \pm 0.007$ | $0.902 \pm 0.007$ |

*Takeaway.* Comparable results with RGIN suggest the framework is robust to encoder choice; the core driver is the self-supervised multi-relation formulation combined with rich graph context.

#### F.6.2 Weight decomposition and vertical stacking

We assessed the effect of *weight decomposition* (basis sharing across relations) and *vertical stacking* (single spmm over a stacked relation matrix) on both efficiency and accuracy. Runtime was measured on the same data and training schedule.

##### Efficiency

Vertical stacking accounts for the dominant speedup versus a naive per-relation pass. Adding weight decomposition maintains this fast regime while reducing parameter count via sharing. Without employing vertical stacking and weight decomposition, MORGaN training exhibits a substantially higher runtime (~seconds per iteration compared to 0.23 seconds per iteration). With vertical stacking but without weight decomposition, the runtime was approximately 4.26iterations per second.

##### Accuracy

With vertical stacking but *without* weight decomposition, we observed slightly higher metrics; however, given the large efficiency/parameter benefits of decomposition and its regularizing effect, we retain it as the default. Reported means ± s.d. over the same splits:

**Table 12:** Performance with vertical stacking but without weight decomposition.

|  | **AUPR** | **AUROC** | **Accuracy** | **F1** |
| --- | --- | --- | --- | --- |
| Vertical stacking and decomposition | $0.888 \pm 0.004$ | $0.907 \pm 0.005$ | $0.915 \pm 0.005$ | $0.917 \pm 0.004$ |
| Vertical stacking and no decomposition | $0.912 \pm 0.010$ | $0.913 \pm 0.004$ | $0.894 \pm 0.010$ | $0.897 \pm 0.012$ |

*Takeaway.* Vertical stacking delivers the primary runtime gain, while weight decomposition preserves that efficiency, reduces parameters through sharing, and serves as an implicit regularizer; we therefore keep decomposition in MORGaN’s default encoder.

## G Family enrichment analysis

## H Pathway enrichment analysis

**Rationale.** Given a ranked list of genes from MORGaN (high score = predicted drug target), *pathway enrichment* asks: “Do the top-ranked genes cluster in curated biological pathways more than we would expect by chance?” If so, that provides *external validity*: the model is concentrating probability mass on coherent processes (e.g., cell cycle, receptor signaling) rather than on idiosyncratic single genes.

**Pipeline in brief.** We use GSEA (Gene Set Enrichment Analysis) in the “pre-ranked” mode:

1. **Rank genes.** Sort all genes by MORGaN’s prediction score.
2. **Choose gene sets.** Use curated pathway collections (e.g., KEGG, GO). Each set is simply a list of genes that participate in a process.
3. **Enrichment statistic.** For each pathway, GSEA computes a running-sum statistic that increases when a pathway gene is encountered high in the ranking and decreases otherwise. The maximum deviation of this walk is the raw enrichment score.
4. **Normalization and significance.** Scores are **normalized** by gene-set size, yielding the **NES** (Normalized Enrichment Score), which lets large and small pathways be compared. Significance is assessed by permutation to form a null distribution; we report nominal *p*-values (NOM *p*) and multiple-testing–corrected **FDR** *q***-values**.

**Analyzed gene sets.** We run GSEA on two sets of predictions: **(A)** all genes predicted as positive by MORGaN, and **(N)** the subset of *novel* positives with no prior annotation. Tables 14–15 and Fig. 8 summarize the most significant results (FDR *<* 0.05).

**Figure 8:**
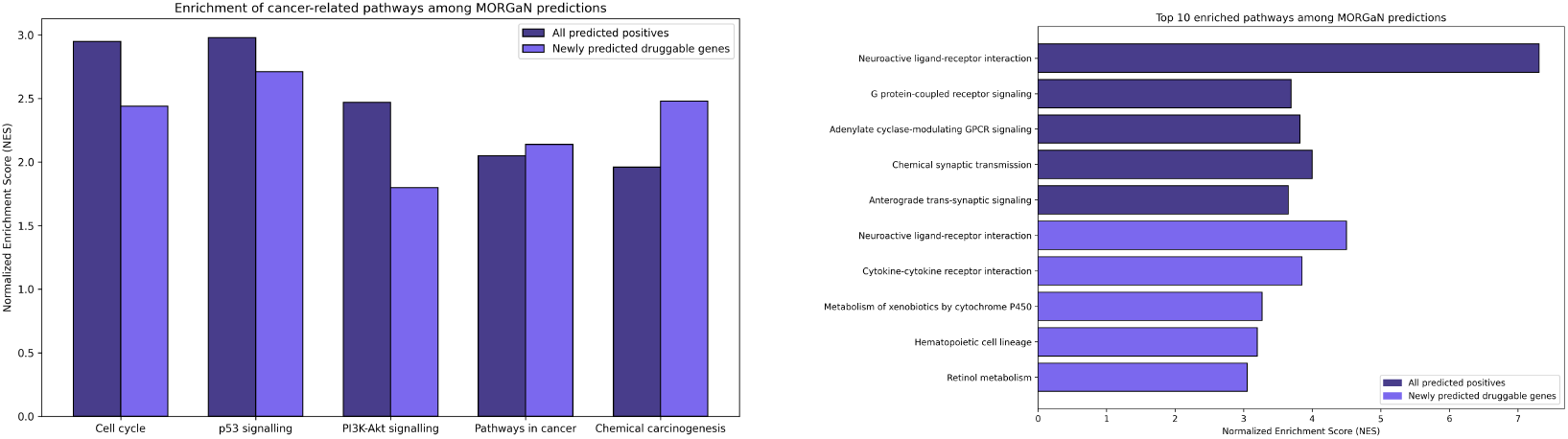
Visual summary of pathway enrichment analysis results. *Left:* Normalized enrichment score (NES) for five hallmark cancer pathways. *Right:* Ten most significant pathways overall. Dark bars = all predicted positives; light bars = novel predictions only.

**Table 13:** Family enrichment among top-tier predictions. Overlap = number of top-tier genes in the family. *p*-values from one-sided Fisher’s exact (greater); FDR = Benjamini–Hochberg.

| Family | Overlap | Odds ratio | $p$-value | Set size | Rank | FDR |
| --- | --- | --- | --- | --- | --- | --- |
| GPCRs | 24 | 4.84 | $3.72 \times 10^{-9}$ | 488 | 1 | $3.72 \times 10^{-8}$ |
| Ion channels | 17 | 5.91 | $3.55 \times 10^{-8}$ | 277 | 2 | $1.77 \times 10^{-7}$ |
| Receptors | 41 | 2.28 | $1.76 \times 10^{-5}$ | 1,769 | 3 | $5.87 \times 10^{-5}$ |
| Cytokine receptors | 5 | 12.50 | $9.54 \times 10^{-5}$ | 39 | 4 | $2.39 \times 10^{-4}$ |
| Transporters | 10 | 1.79 | $6.54 \times 10^{-2}$ | 484 | 5 | $1.31 \times 10^{-1}$ |
| Enzymes | 36 | 1.05 | $4.21 \times 10^{-1}$ | 2,889 | 6 | $7.02 \times 10^{-1}$ |
| Receptor tyrosine kinases | 2 | 1.31 | $4.54 \times 10^{-1}$ | 128 | 7 | $6.49 \times 10^{-1}$ |
| Kinases | 15 | 0.75 | $8.89 \times 10^{-1}$ | 1,627 | 8 | 1.00 |
| Phosphatases | 1 | 0.29 | $9.68 \times 10^{-1}$ | 283 | 9 | 1.00 |
| Nuclear receptors | 0 | 0.00 | 1.00 | 91 | 10 | 1.00 |

**Table 14:** Enrichment of hallmark cancer pathways among MORGaN predictions. Normalized enrichment score (NES), FDR *q*-value and nominal *p*-value (NOM *p*) are shown for both established drug targets (A) and newly predicted candidates (N). All listed pathways pass FDR ≤ 0.05 and NOM *p* ≤ 0.01.

| Pathway | Group | NES | FDR $q$ | NOM $p$ |
| --- | --- | --- | --- | --- |
| Cell cycle (KEGG) | A | 2.95 | 0.00049 | 0.000 |
| Cell cycle (KEGG) | N | 2.44 | 0.00610 | 0.0023 |
| p53 signaling pathway (KEGG) | A | 2.98 | 0.00098 | 0.000 |
| p53 signaling pathway (KEGG) | N | 2.71 | 0.00043 | 0.000 |
| PI3K–Akt signaling pathway (KEGG) | A | 2.47 | 0.00112 | 0.000 |
| PI3K–Akt signaling pathway (KEGG) | N | 1.80 | 0.04700 | 0.0077 |
| Pathways in cancer (KEGG) | A | 2.05 | 0.01580 | 0.0031 |
| Pathways in cancer (KEGG) | N | 2.14 | 0.01050 | 0.0014 |
| Chemical carcinogenesis (KEGG) | A | 1.96 | 0.02410 | 0.0050 |
| Chemical carcinogenesis (KEGG) | N | 2.48 | 0.00057 | 0.000 |

**Table 15:** Top five pathways enriched among all (A) and novel (N) MORGaN-predicted drug targets. Metrics as in. **Table 14**.

| Pathway | Group | NES | FDR $q$ | NOM $p$ |
| --- | --- | --- | --- | --- |
| Neuroactive ligand–receptor interaction (KEGG) | A | 7.31 | 0.000 | 0.000 |
| G protein-coupled receptor signaling (GO) | A | 3.69 | 0.000 | 0.000 |
| Adenylate cyclase-modulating GPCR signaling (GO) | A | 3.82 | 0.000 | 0.000 |
| Chemical synaptic transmission (GO) | A | 4.00 | 0.000 | 0.000 |
| Anterograde trans-synaptic signaling (GO) | A | 3.65 | 0.000 | 0.000 |
| Neuroactive ligand–receptor interaction (KEGG) | N | 4.50 | 0.000 | 0.000 |
| Cytokine–cytokine receptor interaction (KEGG) | N | 3.85 | 0.000 | 0.000 |
| Xenobiotic metabolism by cytochrome P450 (KEGG) | N | 3.27 | 0.000 | 0.000 |
| Hematopoietic cell lineage (KEGG) | N | 3.20 | 0.000 | 0.000 |
| Retinol metabolism (KEGG) | N | 3.05 | 0.000 | 0.000 |

NES measures *how strongly* a pathway is enriched at the top of the ranking after accounting for set size. FDR *q* controls for testing many pathways at once (analogous to a false discovery rate in multiple-hypothesis testing). The bar plots in Fig. 8 compare NES across pathway categories; darker bars refer to results on set (A) and lighter bars refer to set (N).

**Cancer hallmarks.** Both sets recover core oncogenic programs – *cell cycle*, *p53*, *PI3K–Akt*, and composite *pathways in cancer* – indicating that high-scoring genes cluster in well-established cancer biology (Table 14).

**Therapeutically actionable signalling.** The strongest signals are *receptor-mediated* pathways, led by *neuroactive ligand–receptor interaction* and several **GPCR** cascades (Table 15). GPCRs and related receptors are classic drug targets because they are membrane-exposed, ligandable, and already richly represented in approved medicines. Enrichment here suggests MORGaN’s scores align with historically “druggable” target classes rather than random gene families.

**Immune and metabolism niches.** In the novel set (N), we observe *cytokine–cytokine receptor interaction*, *hematopoietic cell lineage*, and xenobiotic/retinol metabolism. These point to *immuno-modulatory* mechanisms (e.g., tuning tumor–immune interactions) and to metabolic processes associated with drug processing and resistance – fertile ground for new targets.

Pathways overlap and are correlated; FDR addresses multiple testing, and NES mitigates gene-set size effects, but some redundancy is expected. Because MORGaN is trained with multi-omic and network context, we consider pathway-level enrichment a complementary sanity check that the model’s global ranking is biologically coherent.

Taken together, the enrichment profile shows that MORGaN both *rediscovers* canonical drug classes (external validity) and *highlights* plausible novel targets for follow-up (novel set N).

## I Local interpretability: case studies

Deep graph models often deliver accurate predictions while leaving the mechanistic “*why*” opaque. We ask: “*Which subgraph structure and which feature dimensions were most influential for MORGaN’s decision on a specific gene?*” Local explanations help users assess faithfulness, spot failure modes, and form testable hypotheses.

To examine MORGaN’s decision process we apply **GNNExplainer** [39], which learns *soft masks* over (i) edges (*M_E_* ∈ [0, 1]^|^*^E^*^|^) and (ii) feature dimensions (*M_F_* ∈ [0, 1]*^d^*). The explainer optimizes these masks to *maximize the mutual information* between the masked inputs and the model’s output for the target node:

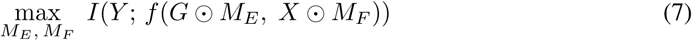

where *f* is the frozen trained model, *G* is the graph (adjacency), *X* are node features, and denotes element-wise masking. In practice, this is implemented with a differentiable surrogate objective (e.g., cross-entropy on the target logit), plus sparsity and entropy regularizers that encourage compact, human-readable explanations. Thresholding *M_E_* yields an *explanation subgraph*; the thicker the edge, the higher its attribution weight.

Fig. 10 displays subgraphs with the top-20 edges by mask weight for four case genes (two established: *EGFR*, *NOTCH1*; two high-confidence novel: *LAMA3*, *IL4R*). The focal node is enlarged; edge width encodes importance. Fig. 9 aggregates the feature mask into a *cancer-type × omic-layer* heat-map, so we can see whether *structure vs. features*, and *which modality*, drove the call.

### I.1 Case studies

**a) EGFR – validating known biology.** The highest-weight edges connect *EGFR* to *TP53*, *CDK2*, and *CTNNB1*. These neighbors sit on well-studied axes that link receptor tyrosine-kinase signaling to proliferation control: *CDK2* is a core cell-cycle kinase (G1/S transition), *TP53* constrains damaged cells from cycling, and *CTNNB1* (*β*-catenin) mediates Wnt pathway transcriptional programs that reinforce growth signals. The feature mask assigns large weights to copy-number and expression channels in lung adenocarcinoma (LUAD) and lung squamous carcinoma (LUSC), indicating that MORGaN’s *per-gene* score for *EGFR* is supported by both (i) a *structural motif* tying EGFR to cell-cycle checkpoints and (ii) *omics evidence* of amplification/over-expression in the histologies where EGFR inhibitors are first-line therapy.
**b) NOTCH1 – pathway-centered evidence.** Instead of a star around *NOTCH1*, the mask emphasizes two tightly connected patterns: (i) a receptor–kinase crosstalk motif involving *ERBB4* and *MAPK9* (JNK), and (ii) a transcriptional decision module with *RBPJ*, the canonical DNA-binding partner for Notch intracellular domain. This says the model is using *multi-hop pathway context* – how Notch signalling routes into MAPK and transcription – rather than just counting direct interactors. Feature-wise, the importance is spread across expression and methylation channels, which is consistent with NOTCH pathway activity being regulated by both ligand/receptor levels and downstream transcriptional state. The selection of small, interconnected motifs implies the predictor relies on *substructures with function*, not just local density or centrality.
**c) LAMA3 – extracellular-matrix lead.** For the unlabeled candidate *LAMA3* (a laminin subunit in basement membrane), salient neighbors include *ITGA4* (integrin receptor) and *SMAD1/2* (TGF-*β* effectors). Together these mark *ECM–integrin–TGF* crosstalk: integrins sense matrix composition and stiffness, transmit signals that modulate SMAD activity, and jointly regulate adhesion, migration, and invasion. The feature mask concentrates in bladder and thyroid contexts, with expression and methylation dimensions carrying the largest weights, suggesting tumor settings where ECM remodeling is particularly informative for the model’s decision. For a novel prediction, a coherent *mechanistic neighborhood* plus aligned feature evidence is stronger than either alone. The model is not “hallucinating” from topology.
**d) IL4R – immune-evasion angle.** The subgraph highlights edges to *AKT2* (PI3K/AKT survival signaling), *TP53BP1* (DNA-damage signaling), and *RAC1* (actin cytoskeleton and motility). This context is expected for *IL4R*, a cytokine receptor that modulates immune and survival pathways: IL-4/IL-13 signaling can activate PI3K/AKT, reshape cytoskeletal dynamics via Rho GTPases, and influence DNA-damage responses indirectly through cell-state changes. The feature mask is strongest in colorectal and lung cancers, with expression and CNA dimensions dominating, again matching settings in which cytokine-driven immune escape and microenvironmental interactions are prominent. Receptor localization (membrane), a signal-integration neighborhood, and high-weight omic channels together form a consistent explanation. Indeed, the explanation aligns with literature linking IL-4/IL-13 signaling to macrophage polarization and immune escape, supporting IL4R as a promising immuno-oncology target.

Overall, the explanations are *compact*, *stable*, and *mechanistically plausible*, letting us trace MOR-GaN’s “YES” decisions back to specific relational motifs and ’omic signals – useful both as a faithfulness check and as a hypothesis generator for downstream experiments.

**Figure 9:**
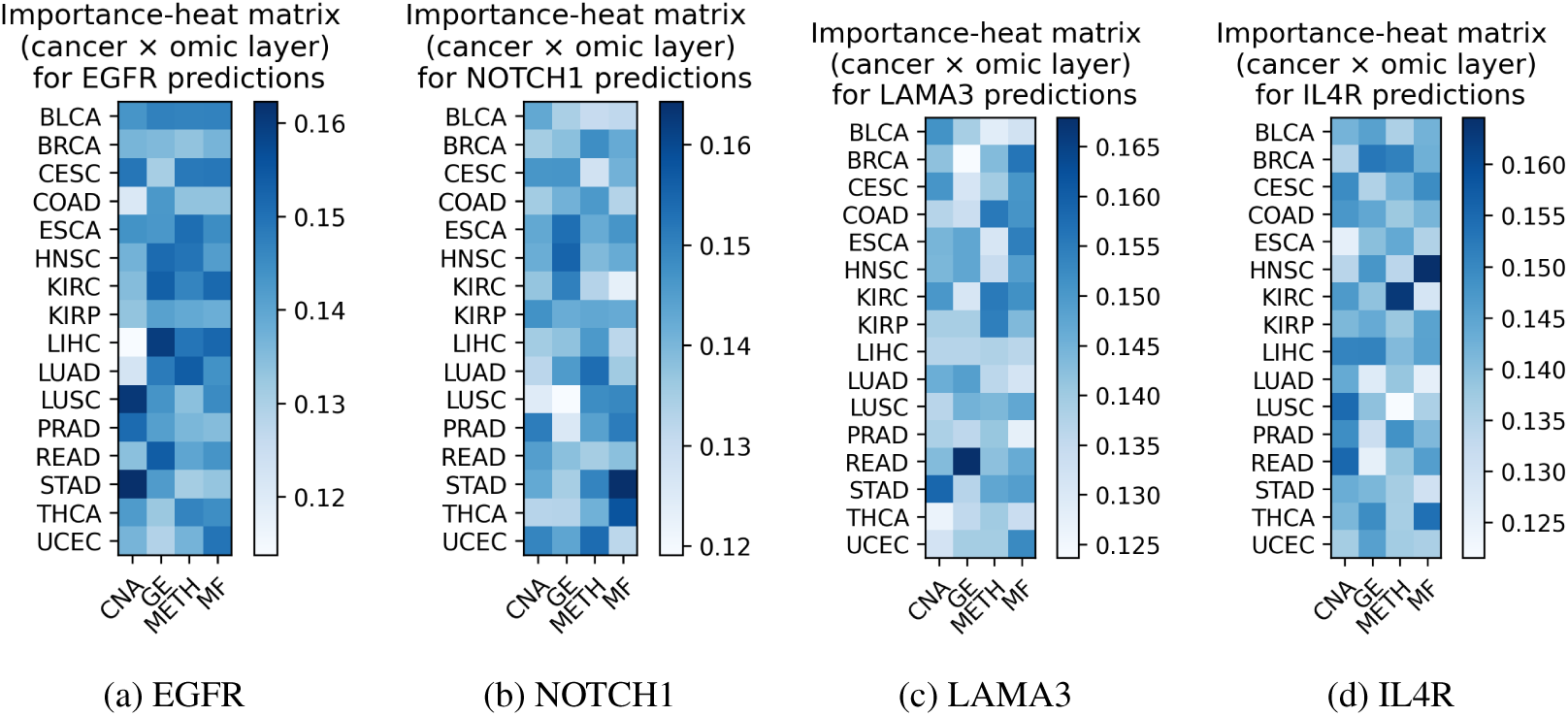
Heat-map visualization of node-feature importance for the same four driver genes. Each panel shows a cancer-type × omic-layer matrix; color intensity is proportional to the contribution weight assigned by GNNExplainer (darker = higher importance).

**Figure 10:**
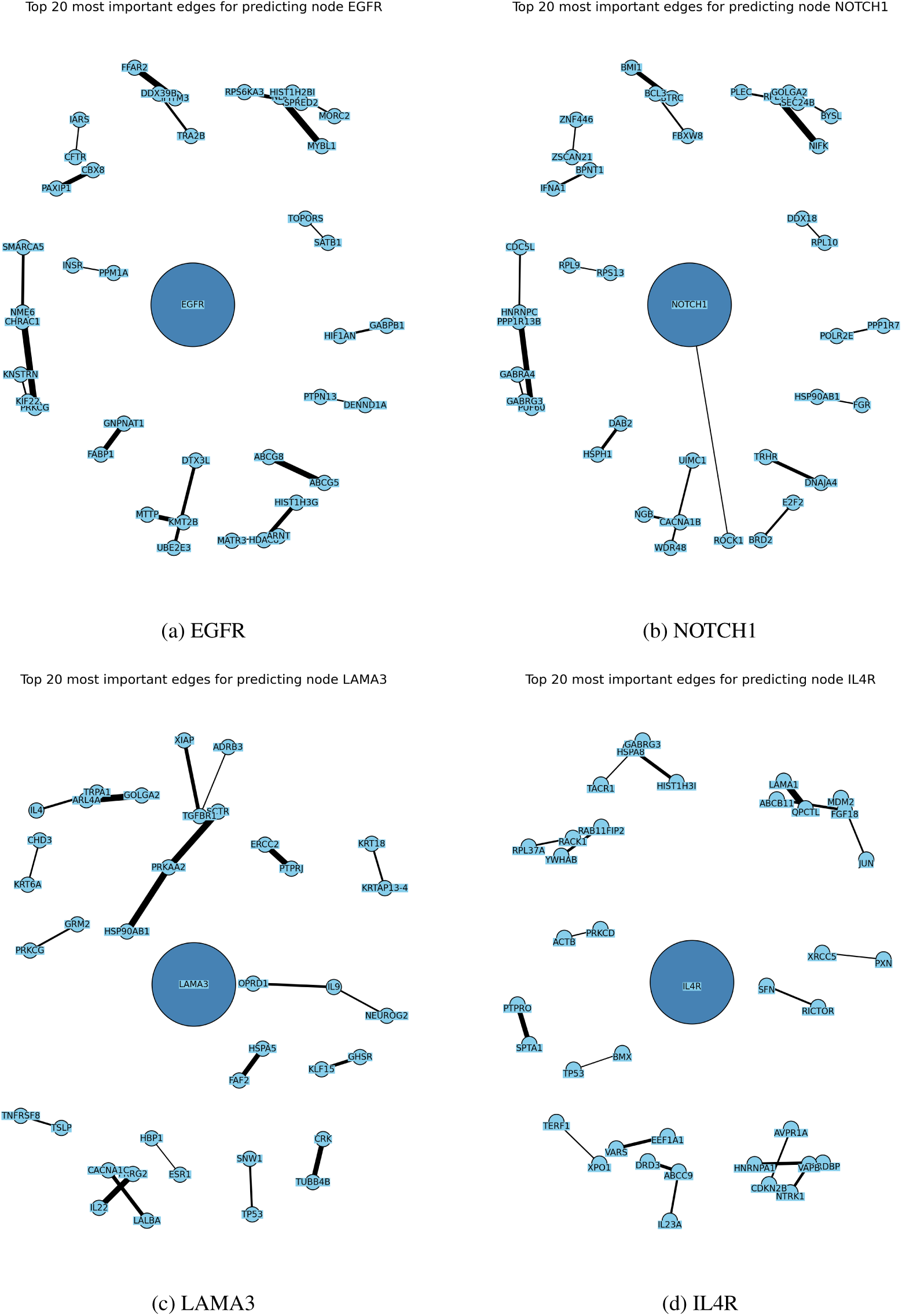
Sub-graphs with the 20 most influential edges (edge width ∝ contribution) for four driver genes. The central node is enlarged and darkened.

## J External concordance

We compared MORGaN’s high-confidence positives (*p >* 0.9) with two external resources: DGIdb [52] and the Finan et al. [4] drug target atlas. Table 16 reports overlaps and proportions. The substantial concordance – particularly the three-way intersection – supports MORGaN’s ability to recover genes independently recognized as drug targets.

Overall, **80.2%** (765/954) of MORGaN’s high-confidence predictions are supported by at least one external resource (DGIdb or Finan), with 63.8% (609/954) shared by both.

**Reproducibility note (MDMNI-DGD).** We attempted to include MDMNI-DGD predictions for a broader comparison; however, the supplementary gene list referenced in their paper was not accessible (the downloadable file appeared corrupted across multiple attempts). We will add this comparison if/when an updated file becomes available.

**Table 16:**
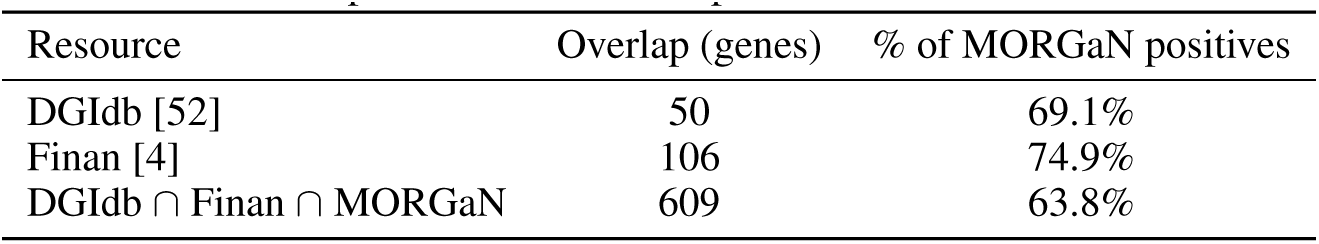
Overlap between MORGaN positives and external resources.

## K Supplementary figures

**Figure 11:**
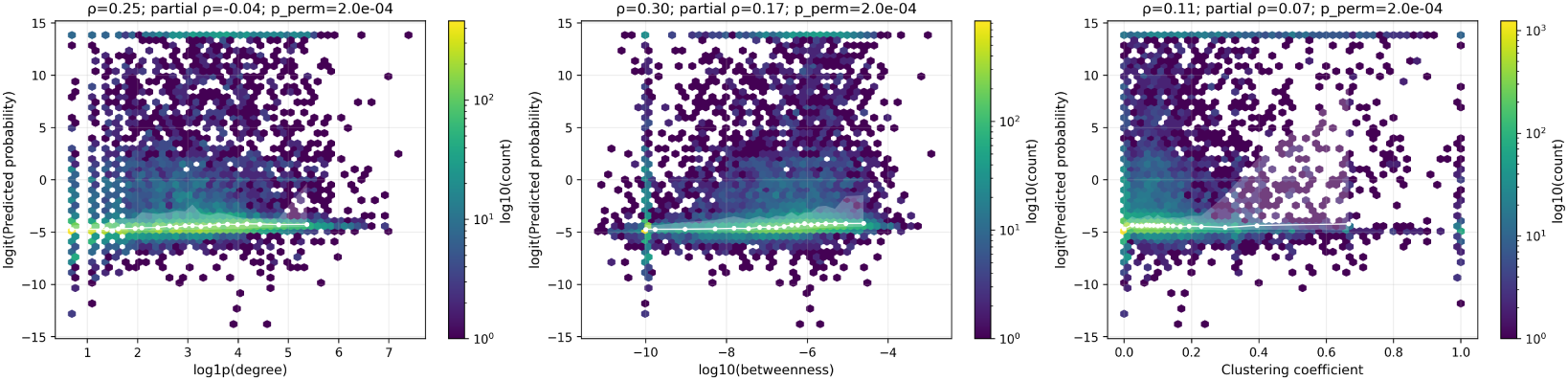
**Hexbin density plots showing the relationship between the prediction scores (logit scale; y-axis) and three network metrics**: degree (log1p, left), betweenness (log10, center), and clustering coefficient (x-axes, right). The white line is a running median. Panel titles report the Spearman correlation (*ρ*), the partial correlation controlling for the other centrality measure, and a permutation p-value. Associations are weak for degree (*ρ* = 0.25; partial *ρ* = −0.04) and modest for betweenness (*ρ* = 0.30; partial *ρ* = 0.17) and clustering (*ρ* = 0.25; partial *ρ* = 0.07), indicating MORGaN is not simply a hub detector but prioritizes genes on information-flow routes and within locally coherent modules.

**Figure 12:**
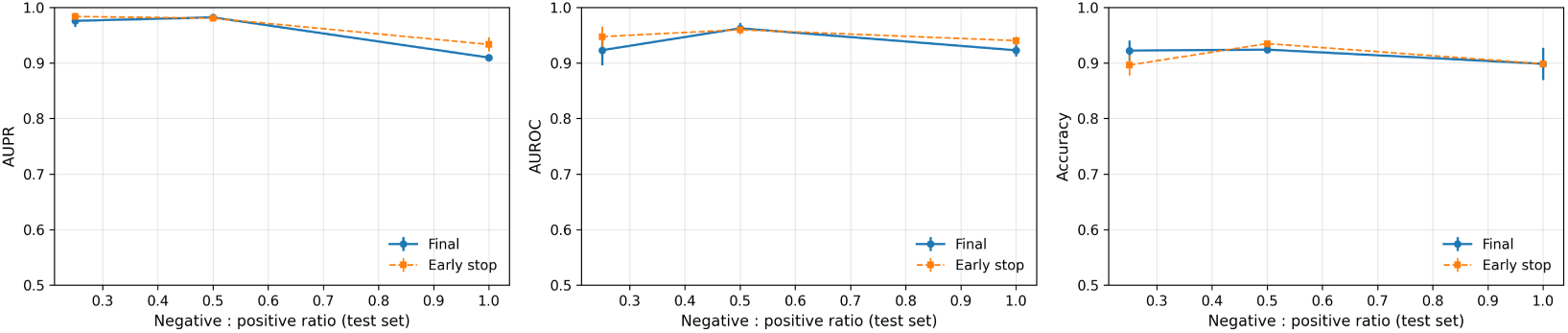
Sensitivity to class balance. Mean ± s.d. across repeated splits for (left) AUPR, (middle) AUROC, and (right) Accuracy as a function of the negative:positive ratio on the test set. Solid lines: final checkpoint; dashed lines: early-stopping checkpoint. AUPR and AUROC remain high across settings; early stopping yields small, consistent improvements at the extremes.

**Figure 13:**
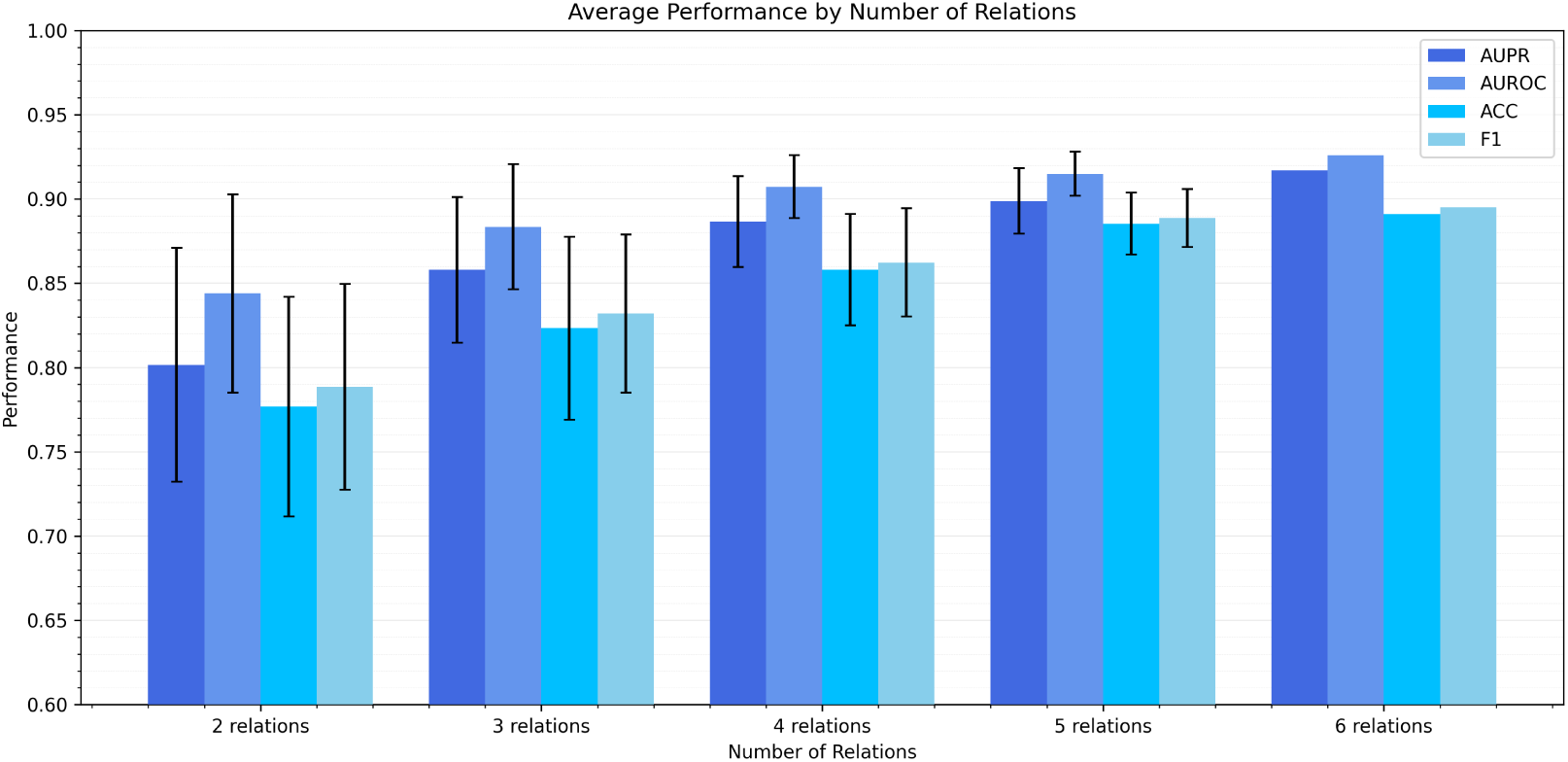
Average performance versus number of relation types. For each *n* = 1 … 6, bars show mean ± s.d. across all combinations of *n* edge types, for AUPR, AUROC, Accuracy, and F1. Performance improves steeply from 1 → 3 relations and then plateaus, indicating additive value from heterogeneous biology with diminishing returns near full coverage.

**Figure 14:**
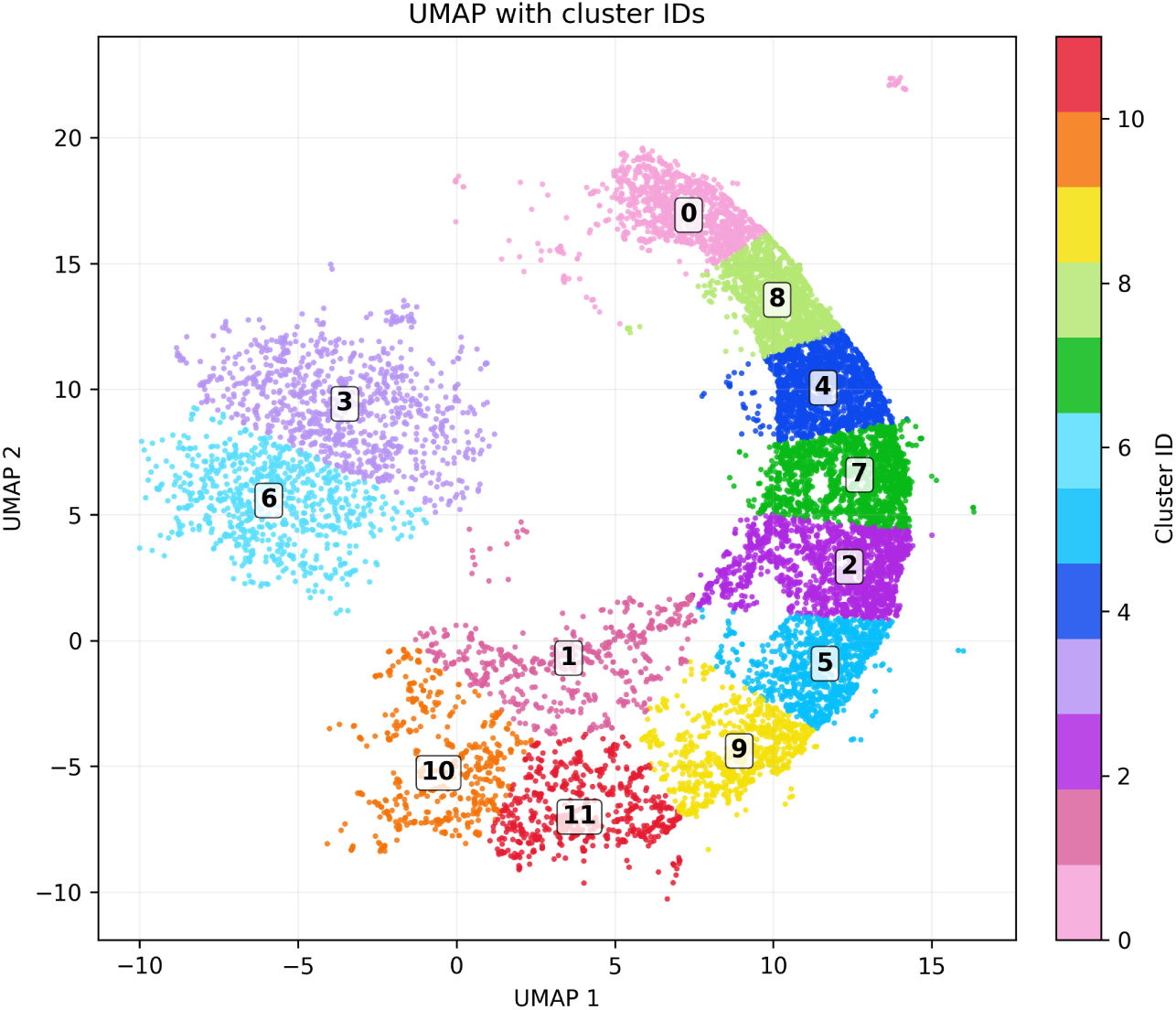
Two-dimensional UMAP of MORGaN node embeddings. (cosine metric, *n*_neighbors_ = 30) colored by k-means clusters (*k* = 12); labels mark cluster IDs. The representation reveals a crescent-shaped manifold of higher-scoring genes and a separate island, providing contiguous segments used for cluster-level pathway enrichment (Fig. 15a–15d).

**Figure 15:**
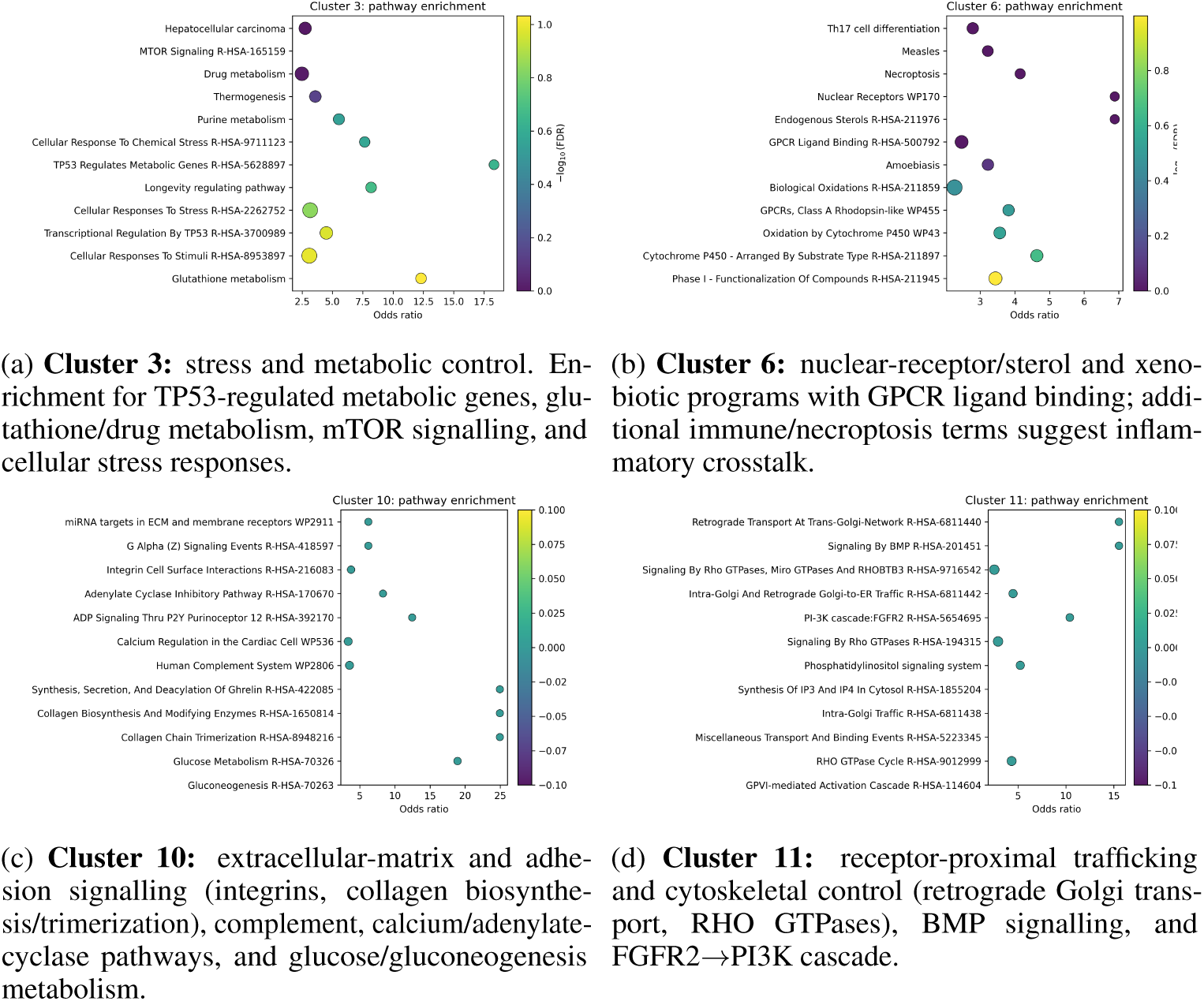
Pathway enrichment of representative UMAP clusters. Bubble plots show the top enriched pathways per cluster (x-axis: odds ratio). Point size is proportional to the overlap (number of cluster genes in the pathway) and colour encodes log_10_(FDR). Distinct clusters capture coherent biological programs spanning metabolism/detoxification, nuclear-receptor signalling, ECM/adhesion and complement, and RTK/RHO-driven signal transduction.

**Figure 16:**
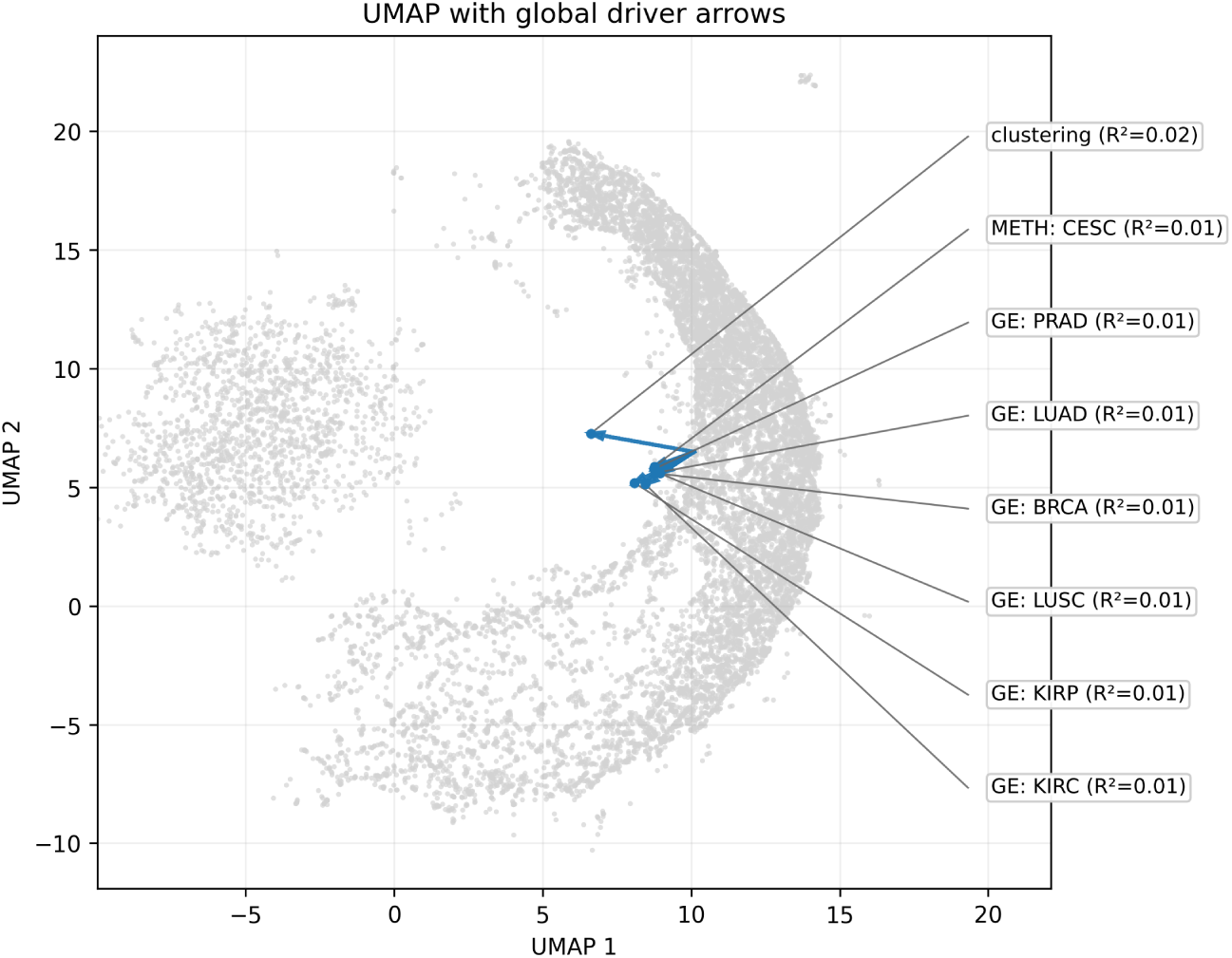
**Envfit projections of external variables onto the centered UMAP coordinates.**Arrows indicate the direction of increasing values; length is proportional to variance explained (reported *R*^2^ for each variable). Cancer-type gene-expression contrasts (e.g., BRCA, LUAD, PRAD, LUSC, KIRC/KIRP), a methylation feature (CESC), and an unsupervised clustering summary align weakly but consistently with the crescent-shaped manifold (*R*^2^ 0.01 0.02), supporting graded, biologically interpretable organization in the embedding.

## L Out-of-distribution experiments

### L.1 Alzheimer’s disease

**Setup.** To test disease-agnostic generalization, we built an Alzheimer’s disease (AD) network using Alzheimer-specific multi-omic profiles (log_2_ fold-change gene expression and chromatin accessibility) and the same six biological relation types used in the pan-cancer graph (derived from [53]). We re-trained MORGaN end-to-end with the identical pre-training and fine-tuning protocol and evaluated on the same split strategy as in the cancer experiments.

**Results.** Performance remains strong under this domain shift, with a small drop relative to oncology (Table 17). This suggests that the self-supervised, multi-relation objective captures disease-general structure that transfers beyond cancer.

**Table 17:** Alzheimer’s disease: mean ± s.d. over splits.

|  | AUPR | AUROC | Accuracy | F1 |
| --- | --- | --- | --- | --- |
| **MORGaN (AD)** | $0.892 \pm 0.022$ | $0.908 \pm 0.009$ | $0.840 \pm 0.009$ | $0.847 \pm 0.008$ |

**Qualitative sanity checks.** Among high-scoring predictions without prior drug targets (“false positives” under our operational binary label), MORGaN prioritizes genes with AD-relevant evidence, including *PDE4D* (amyloid/tau pathology; cognitive decline) [54], *HLA-DRA* (upregulated; neuroinflammation) [55], members of the *HDAC* family (pharmacological modulation ameliorates cognitive deficits in AD models) [56, 57], as well as *NTRK1* (nervous system development) and *NRP1* (neuronal migration, angiogenesis; upregulated in AD models) [58]. These examples support that out-of-distribution predictions remain biologically plausible.

### L.2 Essential genes

**Setup.** To illustrate the task-agnostic utility of MORGaN embeddings, we evaluated a distinct prediction task: gene essentiality. We used proxy labels derived from prior predictions [59] (subset to **E** (essential)) and applied the same training/evaluation protocol (architecture and schedule unchanged), treating this as a separate downstream classification problem.

**Results.** Despite the weaker, prediction-derived labels, performance is competitive (Table 18), indicating that MORGaN learns task-general representations that transfer to essentiality beyond the original objective.

**Table 18:** Essential gene prediction: mean ± s.d. over splits.

|  | AUPR | AUROC | Accuracy | F1 |
| --- | --- | --- | --- | --- |
| **MORGaN (essential)** | $0.765 \pm 0.015$ | $0.835 \pm 0.008$ | $0.772 \pm 0.009$ | $0.797 \pm 0.008$ |

*Takeaway.* Across both experiments, MORGaN’s multi-relation self-supervision yields embeddings that generalize across *diseases* (AD) and *tasks* (essentiality), with only modest degradation under distribution shift and competitive performance under weaker labels.

